# Small Interactor of PKD2 (SIP), a novel PKD2-related single-pass transmembrane protein, is required for proteolytic processing and ciliary import of *Chlamydomonas* PKD2

**DOI:** 10.1101/2023.06.13.544839

**Authors:** Poulomi Das, Betlehem Mekonnen, Rama Alkhofash, Abha Ingle, E. Blair Workman, Alec Feather, Peiwei Liu, Karl F. Lechtreck

**Affiliations:** Department of Cellular Biology, University of Georgia, Athens, GA 30602; Department of Computer Science, University of Georgia, Athens, GA 30602

**Keywords:** TRP channel, cilia, intraflagellar transport

## Abstract

In *Chlamydomonas* cilia, the ciliopathy-relevant TRP channel PKD2 is spatially compartmentalized into a distal region, in which PKD2 binds the axoneme and extracellular mastigonemes, and a smaller proximal region, in which PKD2 is more mobile and lacks mastigonemes. Here, we show that the two PKD2 regions are established early during cilia regeneration and increase in length as cilia elongate. In abnormally long cilia, only the distal region elongated whereas both regions adjusted in length during cilia shortening. In dikaryon rescue experiments, tagged PKD2 rapidly entered the proximal region of PKD2-deficient cilia whereas assembly of the distal region was hindered, suggesting that axonemal docking of PKD2 requires *de novo* ciliary assembly. We identified Small Interactor of PKD2 (SIP), a small PKD2-related protein, as a novel component of the PKD2-mastigoneme complex. In *sip* mutants, stability and proteolytic processing of PKD2 in the cell body were reduced and PKD2-mastigoneme complexes were absent from mutant cilia. Like the *pkd2* and *mst1* mutants, *sip* swims with reduced velocity. Cilia of the *pkd2* mutant beat with normal frequency and bending pattern but were less efficient in moving cells supporting a passive role of the PKD2-SIP-mastigoneme complexes in increasing the effective surface of *Chlamydomonas* cilia.

## Introduction

Cilia and eukaryotic cilia are microtubule-based cell projections with motile and sensory functions. The latter involves channels and receptors in the ciliary membrane, which covers the ciliary axoneme and is continues with the plasma membrane. Rather than being homogenous in composition, the ciliary membrane often contains sub-compartments, in which specific membrane proteins are concentrated. In the auditory cilia of *Drosophila* chordotonal neurons, for example, the TRP channel NompC is localized to parts of the distal zone whereas voltage-gated TRPV channels are present in the proximal zone (Xiang et al., 2022). Also, in Drosophila, the PKD2 orthologue AMO is concentrated near the tip of sperm cilia (Kottgen et al., 2011; Watnick et al., 2003). Similarly, the olfactory cyclic nucleotide-gated channel subunit 1 (OcNC1) is concentrated in the distal segments of rat olfactory cilia (Matsuzaki et al., 1999) and the salt-sensing receptor guanylate cyclase GCY-22 is highly concentrated in the distal region of *C. elegans* primary cilia of ASER neurons (van der Burght et al., 2020). In the latter, the localization of GCY-22 is dynamic requiring motor-driven intraflagellar transport (IFT), a protein shuttle dedicated to the assembly and maintenance of cilia, to continuously capture the receptor along the length of cilia and returning it to the tip by anterograde IFT. Lee et al. reported partitioning of the membrane into actin-dependent corals transiently confining diffusing G-protein coupled receptors (Lee et al., 2018). However, other membrane protein patterns are more static likely involving anchoring of membrane proteins to underlying axonemal structures. Indeed, NompC attaches via its 29 ankyrin repeats to the underlying microtubules, forming a spring-like connection, which contributes to mechanical gating of the channel (Zhang et al., 2015). Finally, some membrane proteins assume additional levels of order around the circumference of cilia, in addition to patterns along the proximo-distal axis. An example is the multiprotein channel complex Catsper, which forms four intricately patterned rows, the race stripes, along the midsegment of mammalian sperm cilia (Chung et al., 2014; Zhao et al., 2022). The observations raise numerous questions including how such membrane proteins are targeted to their specific positions within cilia, how the size of the specialized membrane subdomains is determined and how these regions scale with respect to the overall length of cilia, and how such membrane protein patterns support the cell- and species-specific motile and sensory functions of cilia.

To start addressing these questions, we analyzed the distribution of the TRP channel PKD2 in *Chlamydomonas* cilia, a tractable system for the genetic, biochemical, microscopic and functional analysis of cilia (Dutcher, 1995; Lechtreck, 2016; Pazour et al., 2005; Silflow and Lefebvre, 2001). In the distal two third of the *Chlamydomonas* cilium, PKD2 is attached to the axonemal doublet microtubules (DMTs) 4 and 8 and anchors the mastigonemes, thread-like extracellular polymers of the glycoprotein MST1, to the ciliary membrane (Liu et al., 2020). The proximal third of the cilium is devoid of mastigonemes and PKD2 is more mobile moving by slow diffusion. Here, we analyzed how the PKD2 pattern is established during ciliary assembly and during the repair of PKD2-deficient full-length cilia, and how the pattern adjusts when ciliary length is altered. We also identified Small Interactor of PKD2 (SIP), a novel single-pass transmembrane protein with similarities to the amino-terminus of PKD2. Cilia of the sip mutant largely lack PKD2-mastigoneme complexes and swim with reduced velocity. *Chlamydomonas* PKD2 is cleaved in the cell body and only the two resulting fragments enter the cilia. In *sip* mutants, PKD2 stability and proteolytic processing in the cell body were affected, suggesting a role of SIP in PKD2 processing as a prerequisite for ciliary entry.

## Results

### The PKD2 regions are established early during cilia regeneration

In full-length *Chlamydomonas* cilia, PKD2 is organized into two distinct regions: the proximal region occupying ~1/3 of the cilia, in which PKD2-NG moves by slow diffusion and lacks mastigonemes and the distal region, covering ~2/3 of the cilia, in which PKD2 roughly forms two rows anchoring the mastigonemes to the ciliary surface (Fig. 1A, B) (Liu et al., 2020). The PKD2-NG rows are not always clearly discernable as their distance of ~200 nm is near the limit of resolution of standard light microscopy (DeCaen et al., 2013). The two regions are typically separated by a more or less conspicuous zone lacking PKD2 but for occasional particles passing through, referred to here as the exclusion zone (EZ; Fig. S1A). To study how the PKD2 regions develop while cilia assemble, *pkd2* PKD2-mNG cells were de-ciliated by a pH shock and analyzed at various time points during cilia regeneration using in vivo total internal reflection fluorescence microscopy (TIRFM). A compartmentalization of PKD2-mNG in a proximal and distal region separated by an EZ was apparent in short regenerating cilia with a visible length of ~3 μm (Fig. 1B b-e). Both regions grew as cilia elongated maintaining an average length ratio of ~1:2 between the proximal region and distal region until reaching full length, qualified by the observation that the proximal region was short in cilia with a length of less than 5μm resulting in an average ~1:2.5 ratio (Table S1). Thus, the two PKD2-mNG compartments are established early during cilia regeneration. Remarkably, the data reveal that the PKD2 regions are not established sequentially as cilia grow, i.e., assembling the proximal region first and adding the distal region later once cilia are of sufficient length. Rather the regions adjust in length as cilia elongate, implying that the proximal border of the distal region needs to be moved distally as cilia grow, likely involving the removal of PKD2 anchored in the distal region of shorter cilia.

**Figure 1).**
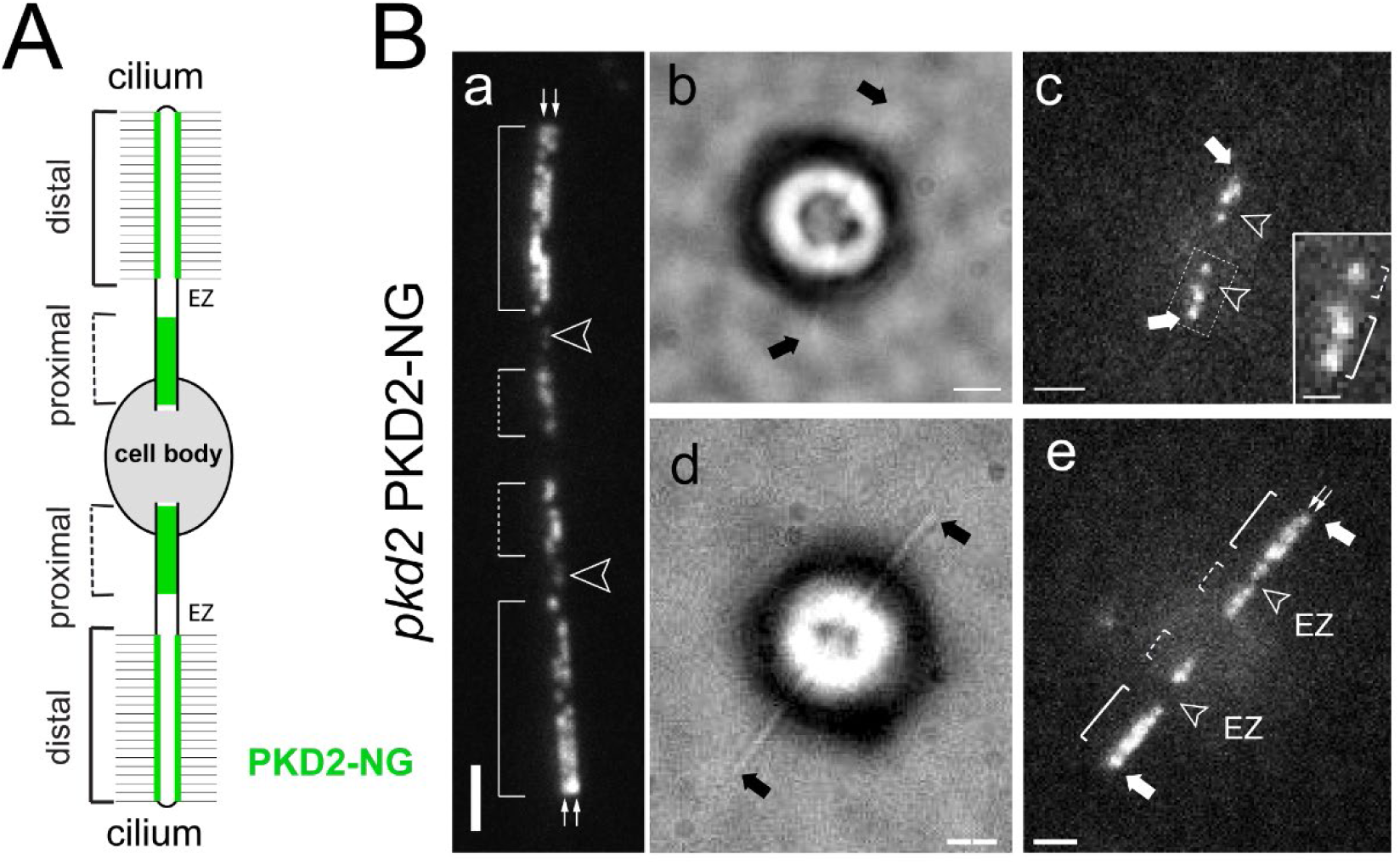
PKD2 regions develop early during ciliogenesis. A) Schematic representation of an adhered *Chlamydomonas* cell with the distribution of PKD2-NG shown in green. B) TIRF images of *pkd2* PKD2–mNG cells with full length (a) and regenerating cilia (b-e). Arrowheads indicate the exclusion zone (EZ) and arrows indicate the ciliary tips. The distal and proximal PKD2-NG regions are marked with brackets. Small double arrows in a point along the two rows of PKD2-NG. Bars = 2μm; bar in inset = 200nm.

### PKD2 regions adjust differently in length in short vs. long cilia

We wondered how the size of the PKD2 regions is affected when ciliary length is perturbed by genetic or pharmacological means. First, we expressed PKD2-GFP in *lf4*, a mutant that assembles cilia exceeding those of control cells 2-3x in length due to the lack of the CDK-related kinase LF4/MOK kinase (Berman et al., 2003). In *lf4 pkd2* PKD2-GFP cilia, the distal stationary region is elongated whereas the size of the proximal region is maintained leading to the length ratio of ~1:4 between the proximal and distal region (Fig. 2A a, b, Table S1). Next, we treated *pkd2* PKD2-mNG for 1 hours with 25mM LiCl, which induces ciliary elongation ((Nakamura et al., 1987; Wilson and Lefebvre, 2004). After LiCl treatment, the average length of cilia was ~16 μm and the average ratio between the distal and proximal PKD2-mNG region had readjusted to ~3.3 (Fig. 2A c, Table S1). The data indicate that in longer than normal cilia the length of the proximal PKD2 region is maintained whereas that of the distal region increases. To determine how the distribution of PKD2 is affected when cilia shorten we treated *pkd2* PKD2-mNG cells with NaPPi to indice cilia resorption (Lefebvre et al., 1978). After a 2h incubation in 20 mM NaPPi, most cells (~90%) had partially resorbed their cilia to an average length of ~5.5 μm (Table S1). Curiously, both the distal and proximal PKD2-NG regions decreased in size maintaining an average ratio of ~2 (Fig. 2A d, e).

**Figure 2).**
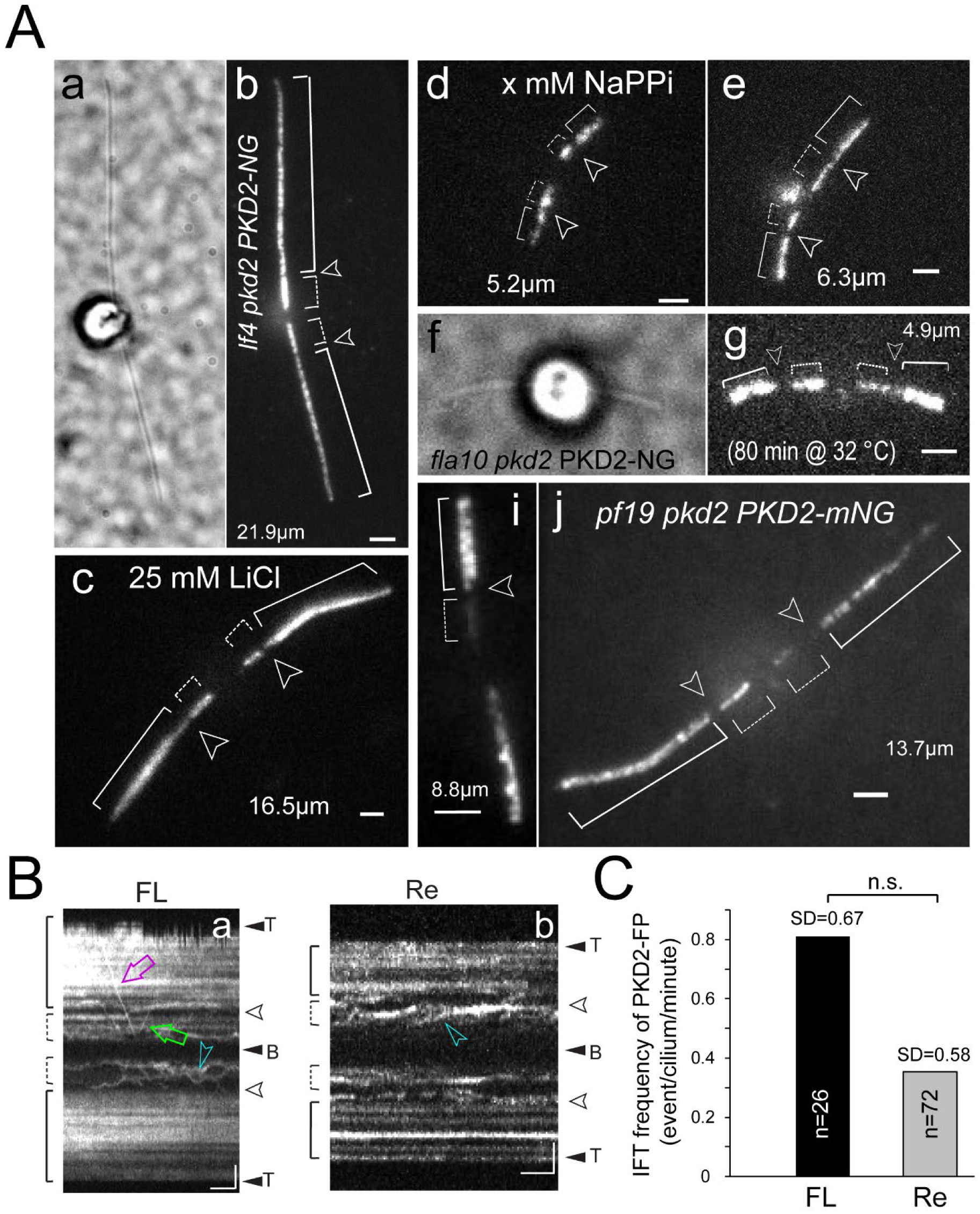
PKD2 regions adjust in size in short cilia and the distal region is extended on long cilia. A) Bright field (a, f) and TIRF images (b-e and g-j) of a *lf4 pkd2 PKD2-GFP* cell (a, b), a cell treated with 25 mM LiCl (c), cells treated with 20 mM NaPPi (d, e), a *fla10 pkd2 PKD2-NG* cell after 80 minutes incubation at 30°C (f, g) and two *pf19 pkd2 PKD2-NG* cells. The ciliary lengths are indicated and the exclusion zones and the distal and proximal regions are marked. Bars = 2μm C) Kymograms of full length (FL; a) and regenerating cilia (Re; b) of a *pkd2* PKD2–mNG cell. The ciliary tips (T), bases (B), the distal and proximal regions, and exclusion zones are marked. Magenta arrow, retrograde IFT; green arrow, anterograde IFT; blue arrowheads, apparent diffusion of PKD2-NG. Bars = 2s and 2μm. D) IFT frequency of PKD2-mNG (anterograde and retrograde combined) in full length (FL) and regenerating (Re) *pkd2 PKD2-mNG* cilia. The SD and number of cilia analyzed (n) are indicated; N.S., difference not significant based on a 2-tailed t-test.

*Chlamydomonas* cilia also shorted in the absence of intraflagellar transport (IFT) (Kozminski et al., 1995). PKD2-NG was expressed in *fla10*, a fast-acting temperature-sensitive allele encoding a motor subunit of the anterograde IFT motor heterotrimeric kinesin-2, which allows one to switch of IFT by a temperature shift (Alfaro et al., 2011; Walther et al., 1994). After incubating *fla10 pkd2 PKD2-NG* cells for 80 minutes at the restrictive temperature of 32°C, cilia had shortened to an average length of ~7 μm but, like observed during NaPPi induced cilia shortening, the ratio between the length of the distal and proximal regions remained at ~2 (Fig. 2A f, g, Table S1).

Since the principal organization of PKD2 in a distal and a proximal region was sustained in the *fla10 pkd2 PKD2-NG* strain at the restrictive temperature, the data also suggest that active IFT is not required to maintain or adjust the size of the PKD2 regions in cilia. Transport of tagged PKD2 by IFT has been previously reported (Huang et al., 2007; Liu et al., 2020). To determine the role of IFT during the assembly of the PKD2-mNG compartments, we compared the frequency by which PKD2-mNG moves by IFT in full length and regenerating *pkd2 PKD2-NG* cilia (Fig 2B, C). In vivo imaging showed similarly low IFT frequencies of PKD2-NG in full-length and regenerating cilia, (0.81 events/min vs 0.35 events/min, respectively; difference not significant; Fig 2C) ((Huang et al., 2007; Liu et al., 2020). While this does not exclude a role for IFT in PKD2 transport, the data suggest that most PKD2-NG enters cilia and organizes into regions in an IFT-independent manner.

The two rows of mastigoneme-associated PKD2 in the distal ciliary region are oriented roughly perpendicular to the plane of ciliary beating, raising the possibility that ciliary motility contributes to the assembly of the rows (Liu et al., 2020). However, in a *pf19 pkd2 PKD2-mNG* strain, which has paralyzed cilia due to the lack of the central pair apparatus, the two rows of PKD2-NG in the distal region and a ratio of ~2 between the distal and proximal PKD2-NG regions was maintained (Fig. 2A i, j, Table S1) (Dymek and Smith, 2012). A subset of *pf19 pkd2* PKD2-mNG cells had longer than normal cilia and, in those cells, the distal-to-proximal ratio was increased, as described for long cilia above (Table S1, Fig. 2A i, j). To summarize, when cilia are abnormally long, only the distal PKD2-NG region was affected and increased in length, whereas both regions adjusted their length in shorter than normal cilia.

### Efficient assembly of distal stationary PKD2 region requires de novo assembly of cilia

To determine if PKD2 can be added in the correct pattern to fully assembled *pkd2* mutant cilia, we mated *pkd2* mutant and *pkd2* PKD2-NG rescue gametes (Fig. 3). After cell fusion, PKD2-NG present in the shared cytoplasm of the zygotes is available for incorporation into the *pkd2*-derived cilia, which initially are devoid of PKD2 (Fig. 3A). In zygotes analyzed ~1h after mixing of the gametes, PKD2-mNG had entered the proximal ~1/3 region of the *pkd2-*derived cilia but only a very few particles were present in the distal region (Fig. 3 B and C). The *pkd2*-derived cilia remained incompletely rescued with PKD2-NG restricted to the proximal region in zygotes analyzed 2 or 3 hours after mixing of the gametes; the distribution of PKD2-mNG in the *pkd2* PKD2-mNG-derived cilia remained apparently unaltered (Fig. 3 B and C). Such incompletely rescued zygotes accounted for 88% of the zygotes summarized from all three time points with the remaining 12% of zygotes having overall weak or no detectable PKD2-mNG signals in the cilia (n = 44 zygotes analyzed; not shown). The latter can be attributed to low PKD2-NG expression in a subset of cells, as it is frequently observed with transgenes in clonal cultures of *Chlamydomonas*.

**Figure 3).**
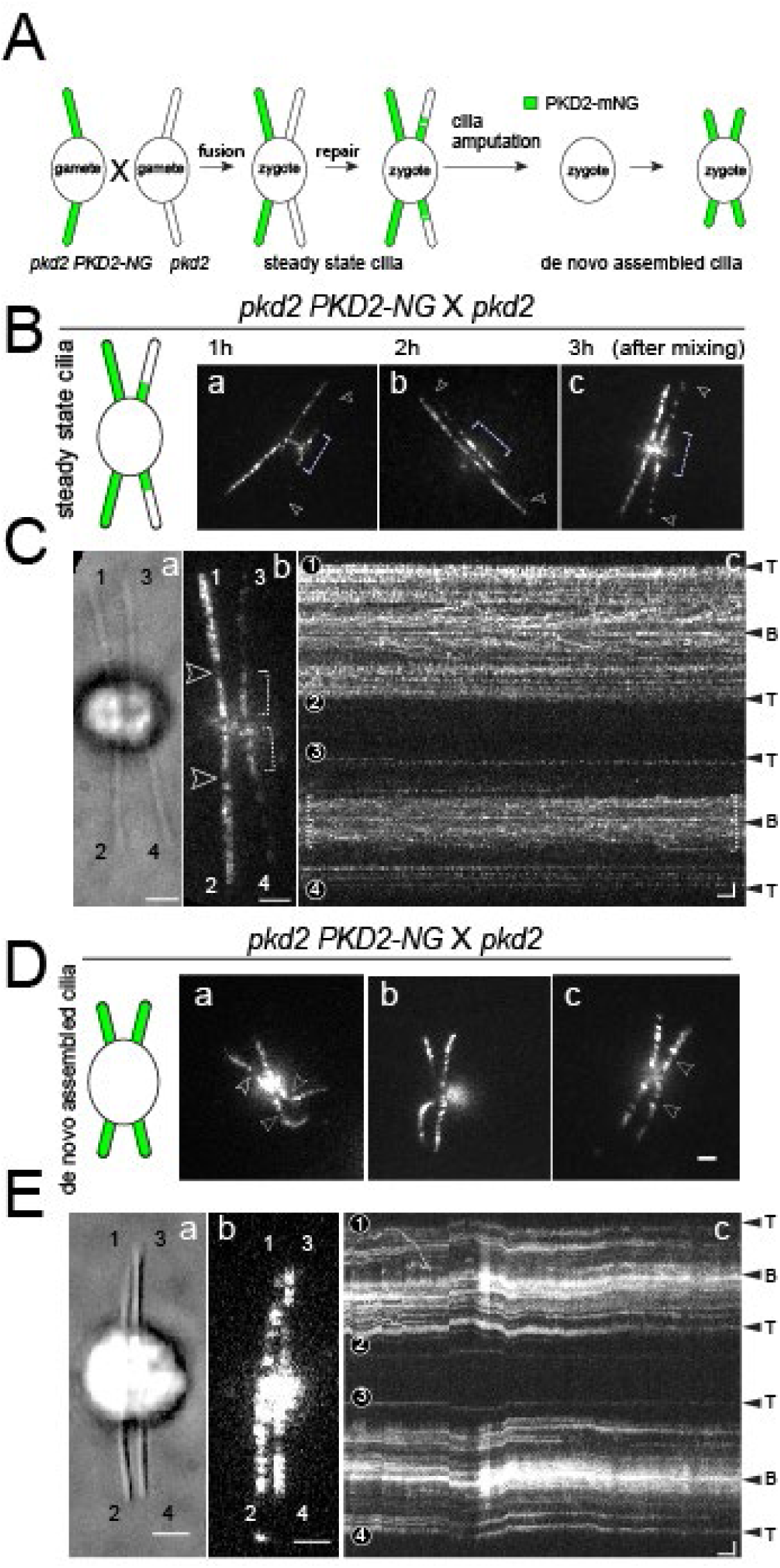
Assembly of PKD2 into the distal region requires de novo cilia. A) Schematic representation of a *Chlamydomonas* mating experiment followed by cilia amputation from the zygote and regeneration. B) Schematic and TIRF images of the zygotes from *pkd2* PKD2-mNG × *pkd2.* Images were recorded at approximately 1h (a), 2h (b), and 3h (c) after mixing of the gametes. Arrowheads indicate the cilia tips. The proximal regions are marked with dotted brackets. Bars = 2μm. C) Bright field (a), TIRF image (b) and corresponding kymogram (c) of a *pkd2* PKD2-mNG × *pkd2* zygote with full-length cilia; the zygote was imaged ~50 minutes after mixing of the gametes. Cilia derived from the *pkd2* PKD2-FP parent are labeled 1 and 2; those derived from the *pkd2* mutant are labeled 3 and 4. The exclusion zone, proximal regions, and ciliary tips and bases are marked. Bars = 2 μm (a, b and a’, b’) and 2 μm 2s (c, c’). D) Schematic and TIRF images of *pkd2* PKD2-mNG × *pkd2* zygotes at ~60 minutes after deciliation of the cells in the mating mixture by a pH shock. Bars = 2μm. E) Bright field (a), TIRF image (b) and corresponding kymogram (c) of a *pkd2* PKD2-mNG × *pkd2* zygote recorded at ~40 minutes after the pH shock. The cilia are labeled 1 to 4; the ciliary tips and bases are marked. Bars = 2 μm (in c for a – c) and 2 μm 2s.

To test if the lack of PKD2 in the distal region is a zygote-specific feature, we deciliated zygotes by a pH shock and allowed them to regenerate all four cilia (Fig. 3A, D, E). In 77% of the 39 zygotes analyzed, the cilia displayed the normal compartmentalization of PKD2-mNG with a 1:2 length ratio of proximal:distal region Fig. 3D, E). The remaining zygotes either largely lacked PKD2-NG or, more frequently, possessed two incompletely rescued and two normal cilia; the latter are likely derived from gametes present in the mating mixture at the time of the pH shock, but only fusing after deciliation and cilia regeneration (not shown).

We also mated *pkd2* PKD2-mNG and wild-type gametes to test the exchange rate of untagged PKD2 in the wild type-derived cilia with PKD2-mNG (Fig. S1B). As described above, PKD2-mNG quickly entered the proximal 1/3 region of the two wild type-derived cilia in 91% of zygotes analyzed but was largely excluded from the distal region of those cilia (Fig S1). The data indicate that PKD2-NG assembly into full-length zygotic cilia is largely limited to the proximal mobile region indicating that PKD2 in this region is more dynamic and in exchange with PKD2 in the cell body. In contrast, assembly of PKD2-NG in the distal region of full-length cilia is incomplete, suggesting that anchoring PKD2-mastigoneme complexes to the axoneme is delayed or inhibited in preexisting cilia.

### Some PKD2-NG is released from the axoneme and cilia during gliding

For *in vivo* imaging, we typically immobilize cells by mixing them 1:1 with immobilization buffer (10 mM Hepes, 5 mM EGTA) to remove extracellular calcium and preventing gliding motility (Bloodgood, 1977; Bloodgood and Salomonsky, 1990). During this form of whole cell locomotion, *Chlamydomonas* cells attach via their cilia to the substrate, here the cover glass, and ciliary transmembrane proteins together with IFT proteins and the retrograde IFT motor IFT dynein/dynein-2 are recruited into ciliary adhesions (Laib et al., 2009; Shih et al., 2013). Stepping of IFT dynein in the adhesions toward the base of the axoneme in the leading cilium drags the cells along the substrate.

To analyze the behavior of PKD2-NG during gliding, we mounted cells in culture medium omitting EGTA. During gliding, a subset of PKD2-NG particles remained stationary causing them to change their relative position within the cilium, moving either toward the base in the leading cilium or the tip of the trailing cilium (Fig. 4 A, B). Such PKD2-NG containing adhesions were typically bright, likely due to a combination of PKD2-NG accumulating in the adhesions and the adhesions being closer to the cover glass surface and therefore generating a brighter TIRF signal (Fig. 4B) (Axelrod, 2003). PKD2-NG often accumulated at the tip of the trailing cilium and the base of the leading cilium of gliding cells. If gliding motility continued, PKD2-NG was occasionally shed from the ciliary tip of the trailing cilium and from the base of the leading cilium leaving trails on the cover glass and in extreme cases depleting most of the PKD2-NG from cilia (Fig. 4B, C; Movie S1). We did not observe obvious differences in the gliding behavior of control and *pkd2* mutant cells, arguing against a role of PKD2 or mastigonemes in gliding motility (Fig. S1 C-E). However, gliding motion likely executes mechanical force on the ciliary membrane and the data suggest that PKD2 responds to such forces and detaches from the axoneme. Release of PKD2 in vesicles from the tip and base of cilia was previously described in *C. elegans* (Wang et al., 2020). Our data document a similar process in *Chlamydomonas* albeit the specificity and significance of PKD2 mobilization and release from cilia are unclear.

**Figure 4).**
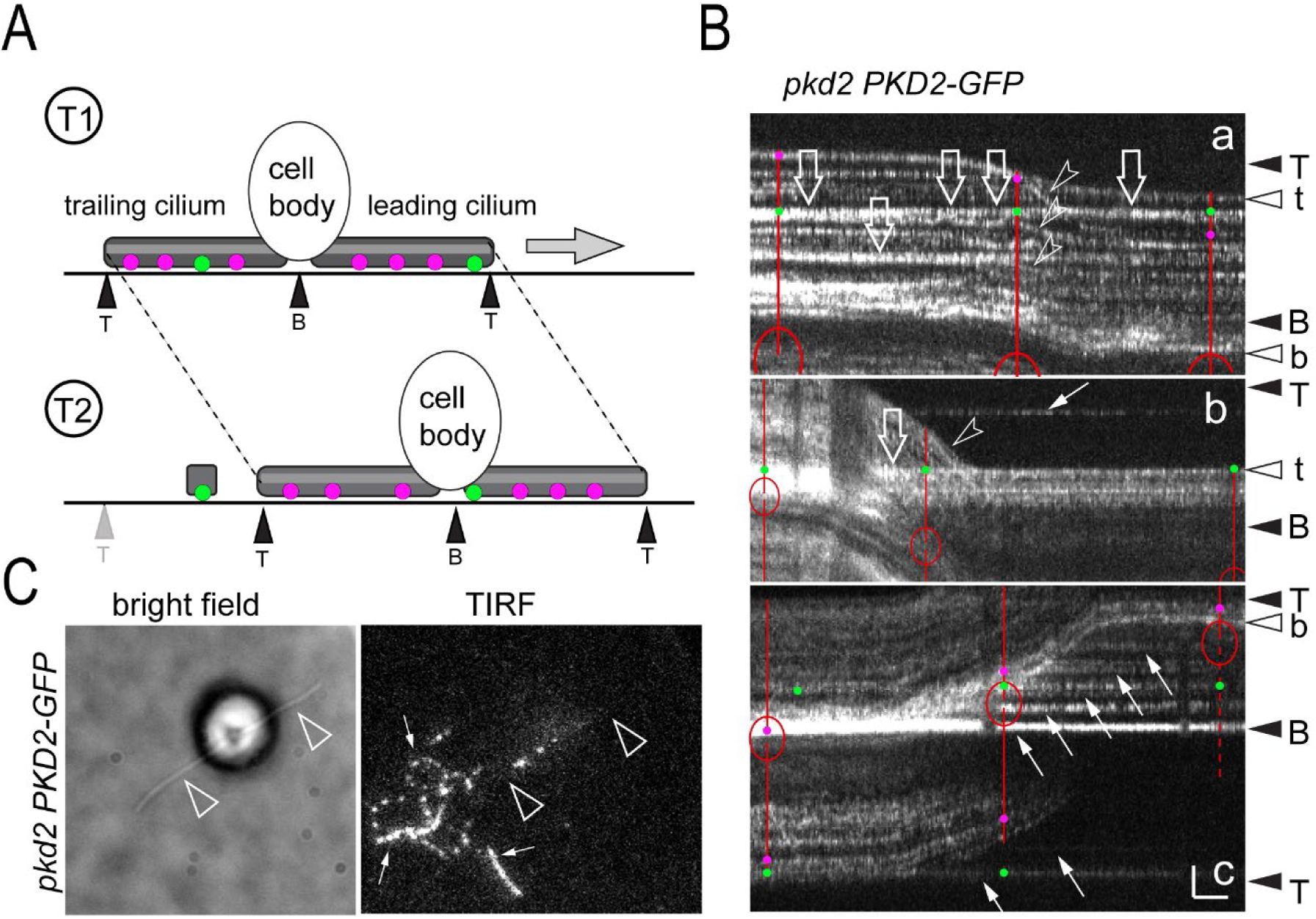
PKD2-NG is mobilized and released from cilia during cell gliding. A) Schematic presentation of *Chlamydomonas* gliding motility. Axoneme-anchored PKD2-NG is indicated by magenta dots; PKD2-NG release from the axoneme and remaining stationary during gliding is indicated by green dots. Occasionally, PKD2-NG is release from cilia during gliding. B) Gallery of kymograms of gliding *pkd2* PKD2-mNG cells. The red lines mark the position of the cell body and cilia at different time points; magenta and green dots, axoneme anchored (i.e., moving with the gliding cells) and stationary (i.e., substrate anchored) PKD2-NG. Open arrows, ciliary PKD2-NG in stationary adhesions; small arrows, PKD2-NG released from the cells at the ciliary tip (b) and base (c); kymogram b and c show the same cell. T and B, ciliary tips and bases prior to gliding movement; t and b, ciliary tips and bases after gliding. Bars = 2 μm 2 s.

### Identification of Small Interactor of PKD2 (SIP) as a novel PKD2-associated protein

To identify additional components of the ciliary PKD2-MST1 complex, we immunopurified PKD2-NG from detergent extracts of *pkd2 PKD2-NG* and *mst1 pkd2 PKD2-NG* cilia using an anti-NG nanobody trap (Fig. S2A). The latter strain was chosen because the lack of MST1 reduces the presence of PKD2-NG in the distal region and proteins specific for the distal PKD2-NG region could be also reduced in this strain compared to the *pkd2 PKD2-NG* rescue strain (Fig. S2B) (Liu et al., 2020). The untagged wild-type strain g1 was used as a control. Silver staining identified several bands in the eluates of the PKD2-NG expressing strains that were absent in the control eluates (Fig. S2A). A prominent band of ~250 kD was present in the *pkd2 PKD2-NG* eluate but not in that of the *mst1 pkd2 PKD2-NG* and control strains and likely represents MST1. The analysis was carried out in several biological replicates (four for the *pkd2 PKD2-NG* and three for the *mst1 pkd2 PKD2-NG* and control strain, respectively) and the eluates were subjected to mass spectrometry (Table S2). Certain abundant (e.g., tubulin) and sticky proteins (e.g., FMG-1) were detected in all 10 samples; PKD2 was detected in the seven experimental samples but not in controls, and, as expected, MST1 was only present in the samples from the *pkd2 PKD2-NG* rescue strain. An additional 35 proteins were identified in most of the *pkd2 PKD2-NG* and/or *mst1 pkd2 PKD2-NG* samples (Table S2). While only present in the experimental samples, we noticed that this list also encompassed proteins that we repeatedly detected in GFP and NG immune-precipitates but not the untagged controls in unrelated experiments; also, the list contained proteins, such as chlorophyll-binding proteins, which we consider unlikely to be associated to PKD2. To triage, we searched for proteins enriched in the *pkd2 PKD2-NG* samples in comparison to the *mst1 pkd2 PKD2-NG* samples. The sole protein fulfilling this criterion was the uncharacterized protein A8JFQ9_CHLRE encoded by the gene CHLRE_11g475150 on chromosome 11 (Fig. S2C; Table S2).

This protein consists of 361 residues, possesses a single transmembrane domain, and is annotated as “Similar to PKD2” in the Phytozome database because it shares similarity with the N-terminal region of *Chlamydomonas* PKD2 on chromosome 17, including a stretch of 30 residues with 80% identity (Fig. S3A). In NCBI blastp searches of *Chlamydomonas* proteins, PKD2 and A8JFQ9_CHLRE identified each other as reciprocal second-best hits (E value 7e-30). In summary, immunoprecipitation and mass spectrometry identified a small protein with certain similarities to PKD2 as a potential interactor of PKD2. For reasons of simplicity, we will refer to A8JFQ9_CHLRE as Small Interactor of PKD2 (SIP). The gene is present in the genomes of various green alga (i.e., Chlorophyta) including Chlamydomodales with MST1-based mastigonemes such as *Gonium pectorale* or *Volvox carteri* as well as species without MST1, such as *Trebuxia sp.,* and *Micromonas sp*., and species that apparently lack the ability to form cilia *(e.g., Scenedesmus sp.)*. Homologues of *Chlamydomonas* SIP were not detected outside of green algae.

To further explore the connection between SIP and PKD2, we obtained strain LMJ.RY0402.143879 from the *Chlamydomonas* CLiP mutant collection; the strain carries an insertion in the second intron of CHLRE_11g475150 (Fig. S2C). A polyclonal antibody raised against recombinant SIP identified a band of ~ 36 in western blots of isolated control cilia, which is close to the predicted molecular weight of SIP of 39,718 (Figs. 5A, S3B). The immunoreactive protein was absent in cilia samples of strain LMJ.RY0402.143879, revealing that this mutant lacks SIP; we therefor refer to this strain as s*ip* (Figs. 5A, C, S3B). The mutant occasionally regained the ability to express SIP, including in mutant progeny obtained by mating; while not further analyzed here, it is likely that cells occasionally gained the ability to remove the transgenic large intron during mRNA processing (not shown). Cilia of control and *sip* equally reacted with anti-SIP in immunofluorescence revealing that this antibody is unsuited for immunocytochemistry (not shown). Whole mount negative stain electron microscopy revealed the lack of mastigonemes from *sip* cilia (Fig. 5B). The absence of mastigonemes was confirmed by immunofluorescence staining with monoclonal anti-MST1 (Nakamura et al., 1996), which further revealed that the pool of MST1/mastigonemes observed in the apical region of control cells is absent in sip mutants cells, as previously described for the *pkd2* mutant (Fig. S3C) (Liu et al., 2020; Nakamura et al., 1996). The strain also swam with reduced velocity as previously described for the *pkd2* and *mst1* mutants (see below).

**Figure 5).**
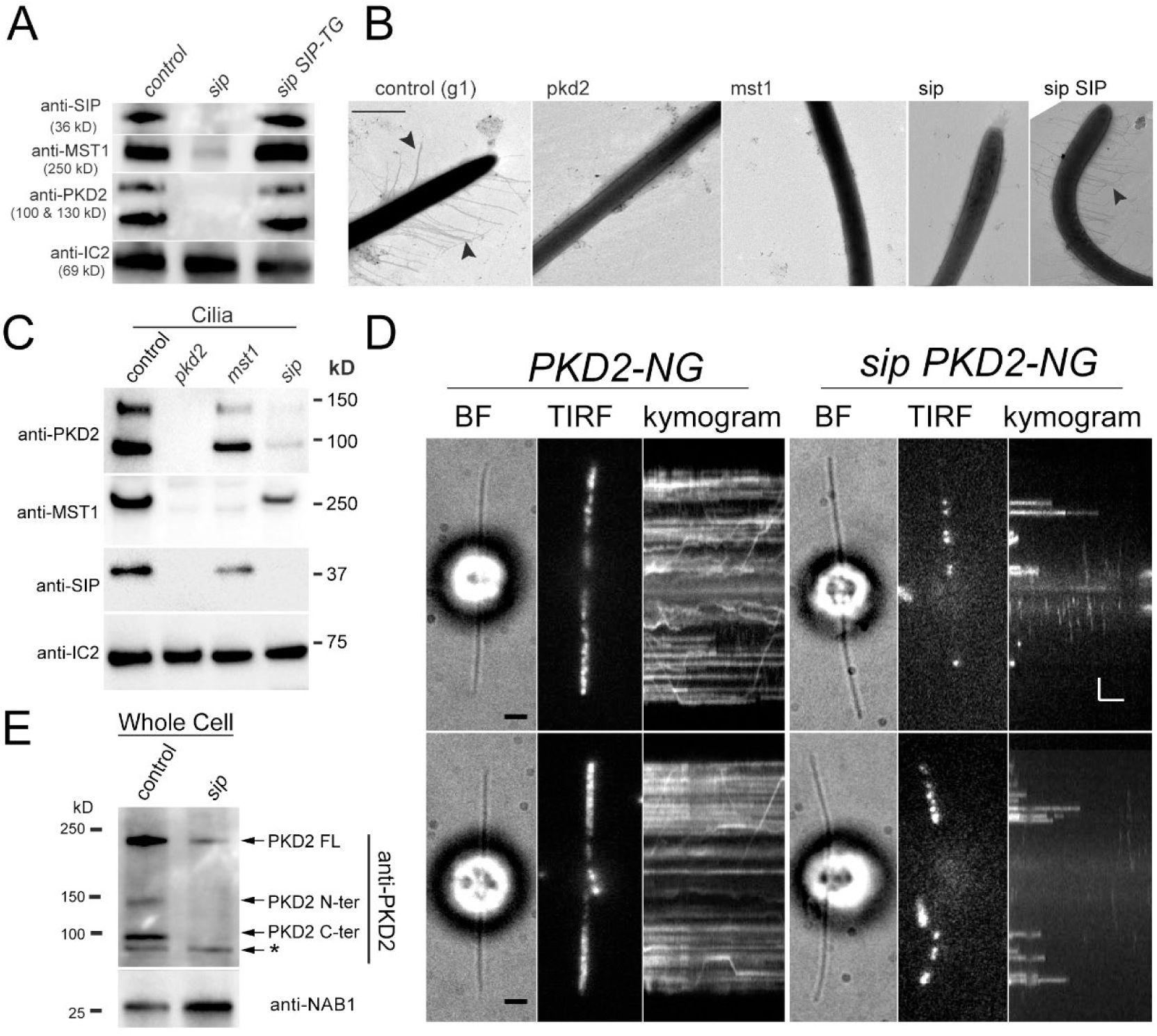
Small interactor of PKD2 (SIP) is part of the PKD2-mastigoneme complex. A) Western blot analysis of cilia isolated from the control, the *sip* mutant and the *sip SIP* rescue strain. The membrane was probed with anti-MINI, anti-PKD2, anti-MST1 and, as a control for equal loading, anti-IC2, a subunit of the outer dynein arm. B) Analysis of cilia from control, *pkd2*, *mst1, sip* mutant, and *sip SIP* rescue cells by whole mount negative stain electron microscopy. Arrowheads, mastigonemes. Bar = 600 nm. C) Western blot analysis of isolated cilia from control and *pkd2*, *mst1* and *sip* mutant cells probed with anti-PKD2, anti-MST1, anti-SIP and, as a control for equal loading, anti-IC2. D) Bright field (BF) and TIRF images and the corresponding kymograms of *PKD2-NG* and *sip PKD2-NG* cells. The proximal PKD2 regions are marked by brackets; open arrows indicate stationary PKD2-NG in the sip PKD2-NG cells; the ciliary tips (T) and bases (B) are indicated. Bars = 2 μm 2s. E) Western blot analysis of whole cell samples of the control and sip strains with anti-PKD2 and NAB1, as a loading control. Note reduction of full-length (FL) and near absence of the N- and C-terminal fragments of PKD2 in the *sip* mutant.

For reasons unknown, our attempts to express SIP fused to GFP failed. Therefore, we expressed untagged SIP in the *sip* mutant; the presence of both the transgenic cDNA-based and the insertional mutant allele was confirmed by PCR (Fig. S2E). Western blotting showed that expression of SIP restored PKD2 and MST1 levels in cilia and electron microscopy showed the presence of mastigonemes on cilia of the *sip SIP* rescue strain (Fig. 5A, B). During cilia fractionation experiments using Triton X-114 phase partitioning, both PKD2 and SIP were present mostly in the soluble matrix fraction with a minor portion remaining attached to the axonemes and only traces detected in the membrane fraction; this behavior sets them apart from dual-lipidated membrane-associated proteins such as carbonic anhydrase 6 (CAH6), which predominately enter the detergent phase during fractionation (Fig. S3E).

Sip was identified by immunoprecipitation of PKD2-NG and, like the *pkd2* mutant, sip mutant cilia lack mastigonemes, a polymer of the extracellular protein MST1. To analyze the interdependence between these three proteins, isolated cilia were from control, *pkd2*, *mst1* and *sip* mutant cells were compared by Western blotting using antibodies directed against PKD2, SIP, and MST1. Since in our hands the monoclonal anti-MST1 failed to detect MST1 on western blots, we raised a novel polyclonal antibody against ~173-residue fragment of MST1 encoded by exon 14; this polyclonal anti-MST1 identified MST1 in Western blot experiments but was unsuited for immunocytochemistry (Fig. S3D). As expected, all three proteins were detected in control cilia with PKD2 running as two bands, the larger N-terminal fragment and the smaller C-terminal fragment, as previously reported (Fig. 5A and C) (Huang et al., 2007). In the *pkd2* mutant, MST1 and SIP were not detected, indicating that PKD2 holds a central role in the complex and is required for the ciliary presence of MST1 and SIP. In *mst1* cilia, PKD2 and SIP were present but significantly reduced (Fig. 5C). PKD2 and MST1 were strongly reduced in *sip* mutant cilia, revealing that SIP is required for the presence of PKD2-mastigoneme complexes in cilia.

To analyze the behavior of residual PKD2 in *sip* cilia, we expressed PKD2-NG in the *sip* mutant and compared PKD2-NG distribution in cilia to that in a control strain expressing PKD2-NG. As expected from the biochemical analysis of cilia, PKD2-NG levels were reduced in sip cilia (Fig. 5 D). However, residual PKD2-NG remained anchored in the distal region of *sip* cilia; PKD2-NG moving by IFT was not observed. Of note, we cannot exclude that residual SIP allowed some PKD2 processing and translocation into cilia.

The near absence of PKD2 from *sip* cilia, raises the question whether PKD2 is trapped in the mutant cell body, which we addressed by western blot analysis of control and mutant whole cell samples. In whole cell samples, anti-PKD2 recognizes full-length PKD2 (~230 kD) and the two proteolytic fragments of 90 (C-terminal fragment) and 140 kD (N-terminal fragment) (Huang et al., 2007; Liu et al., 2020). The overall amount of PKD2 was drastically reduced in *sip* cells and, remarkably, residual PKD2 was mostly uncleaved and the proteolytic fragments were largely undetectable. Previously, proteolytic cleavage of PKD2 was observed in cilia-deficient *Chlamydomonas* mutants indicating that it occurs in the cell body but that only the two fragments enter the cilia (Huang et al., 2007). We conclude that SIP is a novel interactor of *Chlamydomonas* PKD2, required for the stability and apparently proteolytic processing of PKD2 in the cell body as a likely prerequisite for the presence of *Chlamydomonas* PKD2 in cilia.

### The PKD2-mastigoneme complex increases the efficiency of the ciliary beat

The swimming velocity of *sip* mutant cells was reduced by roughly 15%, similar to that of the *pkd2* and *mst1* mutants; expression of transgenic SIP in the sip mutant rescued the motility phenotype (Fig. 6 A and B). To further analyze how PKD2 and its associate proteins MST1 and SIP promotes fast swimming in *Chlamydomonas*, we used high speed video focusing on the *pkd2* mutant. The bending motion of *pkd2* mutant cilia was apparently normal (not shown). However, the beat efficiency, i.e., the distances the cell moved during each beat cycle, of pkd2 cilia was greatly reduced compared to control and pkd2 PKD2-NG rescue cells, explaining the reduced swimming velocity of *pkd2* mutant (Fig. 6 F). This observation supports a role of the PKD2-mastigoneme complex in increasing the effective surface of the cilia, allowing for faster swimming. This concept could also explain the somewhat increased beat frequency of *pkd2* mutant cilia, in comparison to the wild-type and rescue strains as the absence of mastigonemes could reduce the resistance experience by the beating cilia (Fig. 6 E).

**Figure 6).**
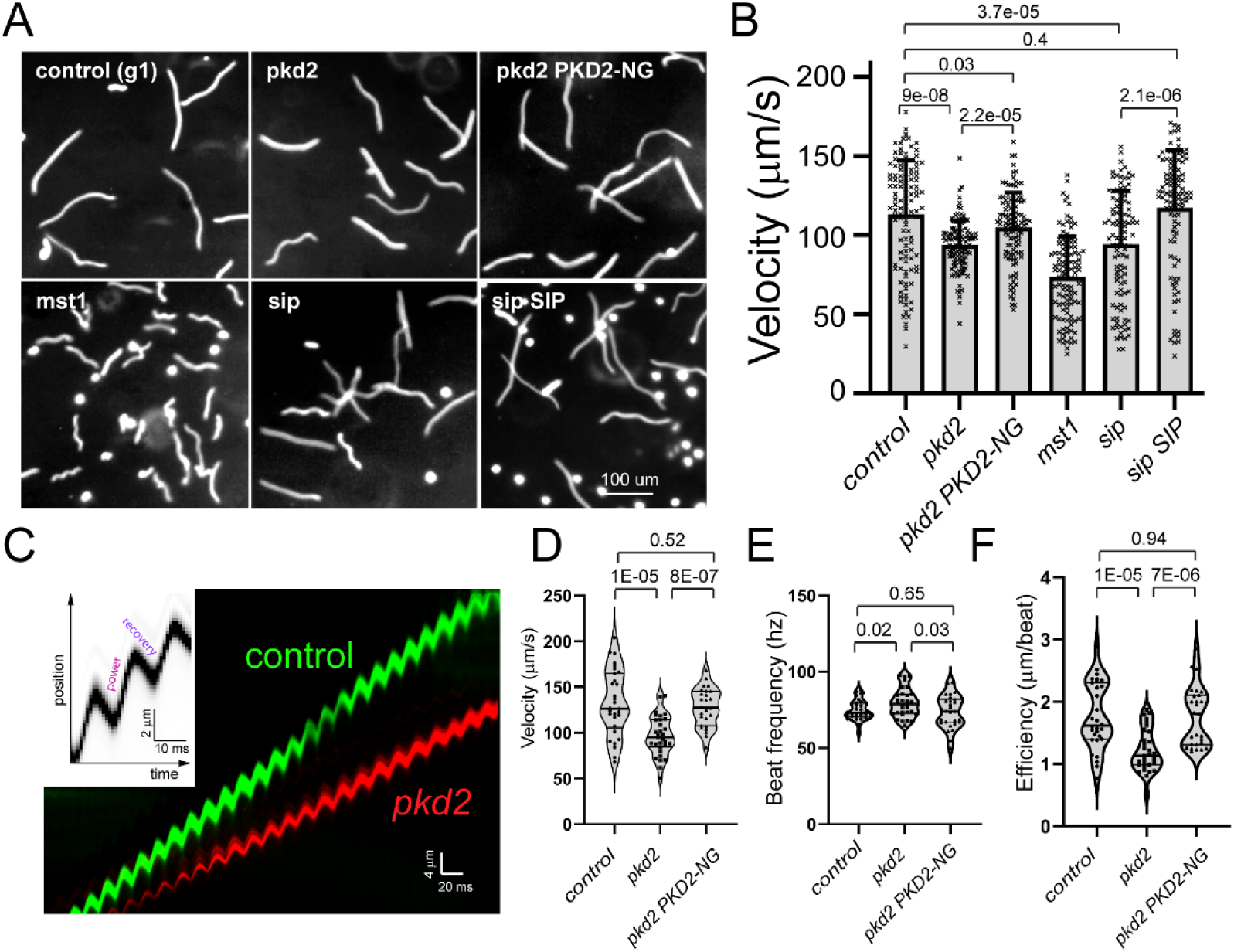
Loss of the PKD2-mastigoneme complex reduced the efficiency of the ciliary beat. A) Micrographs obtained by 1 s exposure showing the swimming paths of control, pkd2, mst1, and sip1 mutant, and pkd2 PKD2-NG and the sip SIP rescue strains. Bar = 100 µm. B) Bar graph of the swimming velocity of various strains. The standard deviations, individual data points and results of a 2-tailed t-test are indicated. C) Overlay kymogram based on high speed recordings of a control (g1; green) and *pkd2* mutant (red) cell. The insert relates a kymogram to the power and recovery stroke. Note that both cells have a similar beat frequency but the pkd2 cell moves with reduced velocity. D) Violin plots of the velocity (a), ciliary beat frequency (b) and beat efficiency (c) of control, pkd2 and *pkd2 PKD2-NG* cells. The results of 2-tailed t-tests are indicated. Shown are the data from one of two experiments each involving 20 or more cells of each strain.

## Discussion

### The size of PKD2 regions is adjusted in a ciliary length-dependent manner

We recently reported that *Chlamydomonas* PKD2 anchors mastigonemes to the ciliary membrane forming two rows of PKD2-mastigoneme complexes along DMTs 4 and 8 in the distal segment of the cilia (Liu et al., 2020). In addition to this radial asymmetry, PKD2 and mastigonemes are also unevenly distributed along the proximo-distal axis of cilia. Whereas PKD2 is mostly stationary in the distal region it often moves by slow diffusion in the proximal region of cilia. The latter region could be comparable to the peri-axonemal inversin compartment in the proximal region of mammalian cilia, which characterized by the presence of inversin and critical for proper ciliary signaling during left-right asymmetry determination and in kidney development (Bennett et al., 2020; Mochizuki et al., 1998; Shiba et al., 2009). In *Chlamydomonas*, the proximal axoneme contains a special subset of inner dynein arms; also, the kinases FA2 and LF5 are restricted to the very proximal end of the ciliary shaft thus above the transition zone (Mahjoub et al., 2002; Tam et al., 2013; Yagi et al., 2009). In addition to these two PKD2 regions in Chlamydomonas cilia, a third region largely devoid of PKD2 separates the proximal and distal PKD2 regions. The overall pattern suggests the presence of distinct PKD2 binding sites in cilia, i.e., stable docking sites on DMTs 4 and 8 and weaker binding sites for PKDs in the proximal region separated by a region devoid of PKD2 binding sites. As with the radial asymmetry of PKD2, we consider yet unknown differences in the underlying axoneme the most likely cause for the proximo-distal differences.

Axonemes grow by addition of tubulin and other proteins to the distal end and proteins that are present in the proximal region of the mature cilia could be delivered and assembled early during cilia formation, whereas those specific for the distal segment will follow later. However, our data show that the borders of the two PKD2 regions shift during cilia assembly as well as during induced shortening. Likely, the reorganization of PKD2 involves changes in the underlying axoneme generating and eliminating sites for stable and transient binding of PKD2. During the assembly of *Drosophila* auditory cilia, dynein motor complexes are initially not confined to their proximal target zone, and ectopic complexes initially observed in the distal zone are later removed (Xiang et al., 2022). Also, OcNC1 is initially present in the proximal segment of assembling rat olfactory cilia and later moved to its final position in the distal segment (Matsuzaki et al., 1999). Thus, the axoneme and membrane are not static along the proximo-distal axis but adjust their composition as cilia grow. This apparently also involves the trimming of proteins incorporated during earlier stages of cilia assembly during cilia maturation. Here, we add that in *Chlamydomonas* such dynamics also occur when the length of cilia is altered by drug treatment or the lack of IFT.

### Assembly of PKD2 into axoneme-anchored rows requires de novo assembly of cilia

In dikaryon experiments, PKD2-NG was restored in the proximal region of the PKD2-deficient mutant-derived cilia whereas its assembly into the distal region was incomplete, typically consisting of just a few stationary PKD2-NG particles. Similarly, exchange of PKD2 with PKD2-NG proceeded rapidly in the proximal region of the wild type-derived cilia of *pkd2 PKD2-NG* × *PKD2* zygotes but was rarely observed in the distal region. The observations reveal that, while addition and exchange PKD2-NG in the proximal region occurs quickly, adding PKD2 to the distal region of fully assembled cilia is a rather slow process, suggesting that efficient axonemal anchoring into two rows requires *de novo* assembly of cilia. Since PKD2-NG enters into zygotic cilia, it is likely that PKD2-NG docking to fully assembled axonemes is impeded because its docking sites on the DMTs 4 and 8 are obscured or absent. Currently, it is unknown whether PKD2 interacts directly with the DMTs or via a linker protein. A putative linker protein or the intracellular regions of PKD2 could integrate into the microtubules during elongation whereas appending PKD2 or its linker to fully formed DMTs could be difficult. Alternatively, PKD2’s axonemal binding sites could be obscured in mature axonemes, e.g., by posttranslational modifications of tubulin.

The PKD2-mastigonemes complexes are targeted with high precision to DMTs 4 and 8, suggesting that docking sites are only present on these two doublets and raising the question how such radial asymmetry is generated. In sperm flagella, distinct patterns of tubulin posttranslational modifications between DMTs have been described raising the possibility that PKD2 docking depends on a specific PTM pattern on DMTs 4 and 8 (Gadadhar et al., 2023). Furthermore, numerous structural specializations typical for one or a subset of the nine DMTs, e.g., B-tubule beaks, the DMT 1-to-2 bridge, absence of ODAs from DTM 1 etc., have been identified in *Chlamydomonas* cilia indicating the presence of biochemical differences between the different doublets (Bui et al., 2012; Dutcher, 2020). The DMTs are continuous with the basal body triplets, each of which possess unique features with respect to their association to basal apparatus fibers and ciliary roots, centrin fibers within the triplet cylinder, and their position within the cell and with respect to the mother basal bodies during their genesis (Geimer and Melkonian, 2004). Thus, it is likely that the distinct features of the axonemal DMTs are related to the individual features of the triplets that nucleate them.

### SIP promotes proteolytic processing and ciliary entry of PKD2

To identify a possible linker protein required for the docking of PKD2 to specific DMTs, we isolated PKD2 complexes from ciliary detergent extracts and identified SIP as single-pass transmembrane protein that interacts with PKD2. Single-pass transmembrane proteins are part of other ciliary channels, for example, the Catsper channel complex in mammalian sperm flagellar. This complex contains at least three single-pass transmembrane proteins (e.g., CatSper γ, CatSper δ, Catsper ζ), which are required for proper trafficking, assembly and/or function of the complex in the midsegment of sperm flagella (Chung et al., 2017; Singh and Rajender, 2015). Further, sodium channels encompass single-pass beta subunits, which regulate channel gating and localization and form links to the underlying cytoskeleton (Isom, 2001).

The overall amount of PKD2 was greatly reduced in *Chlamydomonas sip* mutant cilia, including in the mastigonemes-free proximal segment. Our data suggest that SIP is involved in the processing of PKD2 in the cell body prior to translocation into the cilia. *Chlamydomonas* PKD2 is cleaved in the large extracellular domain between helix 1 and 2 and only the two resulting fragments enter the cilia (Huang et al., 2007). In *sip* cells, the overall amounts of PKD2 was reduced and that the PKD2 fragment were essentially undetectable, indicating a role of SIP in PKD2 stabilization and proteolytic processing. The presence of SIP-encoding genes is limited to the genomes of various green algae including those with *Chlamydomonas*-like mastigonemes but also species, which lack mastigonemes/MST1 and even cilia. This suggests that SIP is a green algal PKD2 processing factor rather than being specifically required for ciliary localization or axonemal binding of PKD2. PKD2 docking might involve a component, which tightly associated with DTMs 4 and 8 and therefore unlikely to be released by detergent treatment as used here to identify SIP. Future work using proximity labeling techniques using PKD2- or SIP-BirA fusions might provide a better strategy to shed light on the axonemal docking site of PKD2.

## Materials and Methods

### Strains, culture conditions and genotyping

*Chlamydomonas* strains used in this study are enlisted in Table 2 and they are wild type (CC-620), g1, *pkd2, lf4* (CC-4534)*, pkd2* PKD2-mNG, *pkd2* PKD2-GFP, *lf4 pkd2* PKD2-GFP, *sip* (<u>LMJ.RY0402.143879</u>), *sip SIP*, *mst1* (LMJ.RY0402.052413) and *mst1 pkd2* PKD2-mNG. The *lf4 pkd2* PKD2-GFP strain was generated by mating *lf4* and *pkd2* PKD2-GFP; progeny was analyzed by PCR, phenotyping ad TIRF microscopy. Information regarding *pkd2,* PKD2-FP rescued strains, *mst1 and mst1 pkd2* PKD2-mNG were previously reported *(Liu et al., 2020)*. The *pkd2* (LMJ.RY0402.204581), *sip* (<u>LMJ.RY0402.143879</u>) and *mst1* (LMJ.RY0402.052413) were acquired from the *Chlamydomonas* Library Project (CLiP; https://www.chlamylibrary.org/allMutants) (Li et al., 2019). Cells were grown in modified minimal (M) medium (https://www.chlamycollection.org/methods/media-recipes/minimal-or-m-medium-and-derivatives-sager-granick/) and maintained at 22°C with a light/dark cycle of 14:10 h; large cultures used for cilia isolation were aerated with air enriched with 0.5% CO_2_.

**Table 1).**
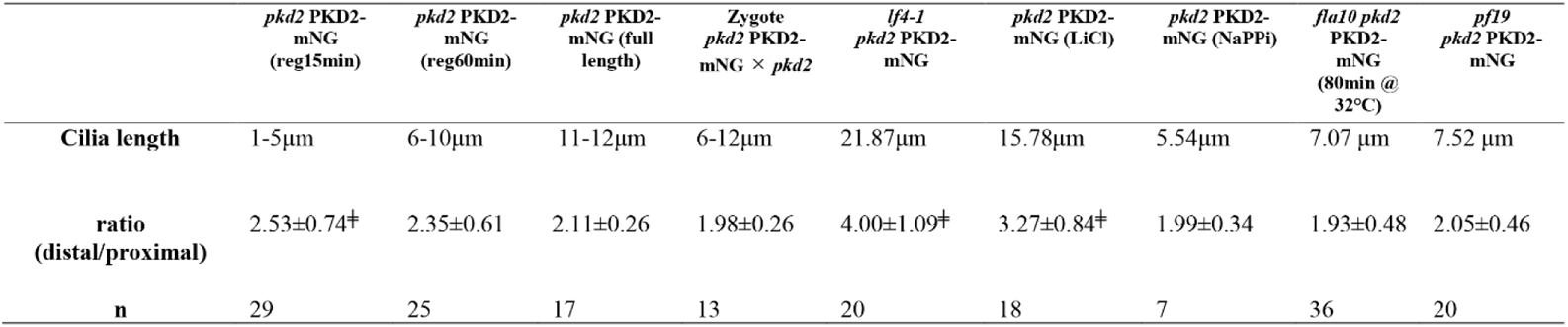
Length ratio of proximal and distal region of PKD2-mNG. N = cilia analyzed. ^╪^, P<0.05 in two tailed t-test compared to wt (full length).

**Table 2).** Strains used in this study. TG, transgenic strain

Table 1) Length ratio of proximal and distal region of PKD2-mNG N = cilia analyzed. <sup>‡</sup>, P<0.05 in two tailed t-test compared to wt (full length).
| Name | Genotype | Reference |
| --- | --- | --- |
| CC-620 (Wild-type) | *Nit, nit2, mt<sup>+</sup>* | *Chlamydomonas* Genetics Centre |
| *gl* (wild-type) | *nit1, agg1, mt<sup>+</sup>* | (Pazour et al., 1995) |
| *pkd2* (progeny selected from *pkd2*<sup>CLiP</sup> backcrossed twice with *gl*) | *cw15, pkd2, mt<sup>-</sup>* | (Liu et al., 2020) |
| *lf4* (CC-4768) | *lf4, mt<sup>+</sup>* | *Chlamydomonas* stock centre |
| *pkd2* PKD2-mNG-TG + *HYG* | *pkd2, mt<sup>-</sup>* | This study |
| PKD2-GFP-TG | *mt<sup>+</sup>* | (Huang et al., 2007) |
| *lf4* *pkd2* PKD2-GFP-TG |  | This study |
| *sip*<sup>CLiP</sup> | *cw15, mini, mt<sup>-</sup>* | This study; (Li et al., 2019) |
| *Sip* SIP-TG | *mt<sup>-</sup>* | This study; (Li et al., 2019) |
| *mst1* (LMJ.RY0402.052413) | *cw15, mst1, mt<sup>-</sup>* | (Li et al., 2019; Liu et al., 2020) |
| *mst1* *pkd2* PKD2-mNG-TG | *mst1, pkd2* | (Liu et al., 2020) |

PCR analysis of isolated genomic DNA was used to confirm the presence of inserted cassette in the mutant. Primers 4 and 5 (Table 3) were used to track the *sip* insertional allele, primers 8 and 9 were used to amplify a part of the g*β* gene to verify DNA quality (Zamora et al., 2004).

**Table 3).**
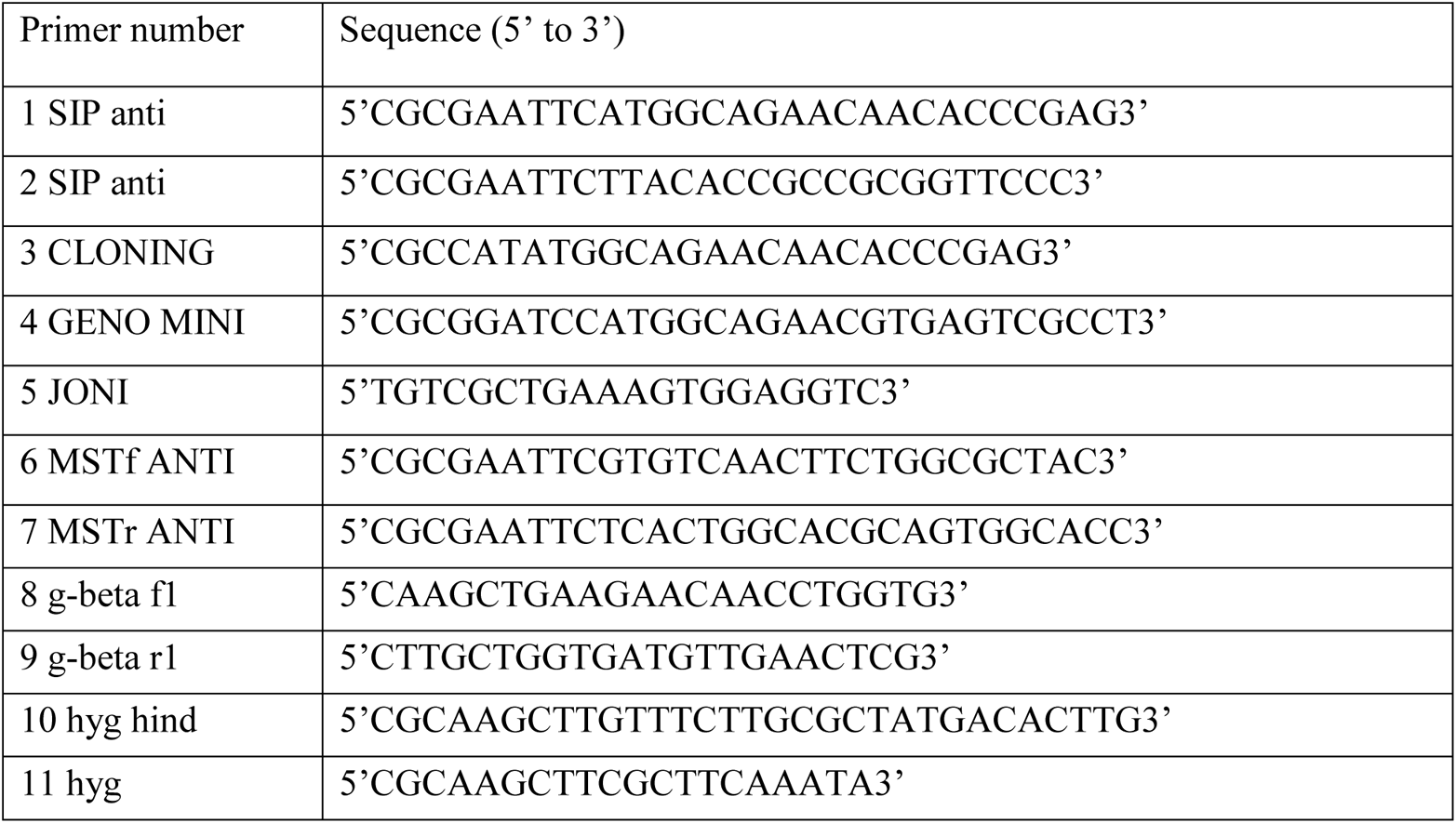
Primers used in this study.

### Transgenic strain generation

To express SIP in the *sip* mutant, cDNA of *SIP* gene was amplified from using primers 2 and 3 (Table 3) and *Chlamydomonas* cDNA as a template. To provide antibiotic resistance the hygromycin cassette was amplified using primer 10 and 11 (Table 3). Amplified *SIP* cDNA and the hygromycin (Hyg) cassette were inserted into the pGenD vector resulting into the formation of pGenD-SIP+Hyg. The resulting plasmid was linearized using XbaI and transformed in the *sip* mutant by electroporation (Invitrogen Neon^TM^ Transfection system). Transformants were selected on TAP plates containing 20µg/ml hygromycin (Bio Basic). Clones expressing SIP protein were identified using whole mount transmission electron microscopy based on the restoration of mastigoneme rows on the ciliary surface. The presence of SIP, PKD2 and MST1 in cilia was confirmed by western blot analysis and the *sip SIP* genotype was confirmed using PCR. 1 out of more than 60 transgenic clones expressed SIP protein.

### Ciliary regeneration

Swimming vegetative cells or gametes were deciliated by a pH-shock in M medium, transferred to fresh M medium, and allowed to regrow cilia under constant light with agitation for ~50min before transferred to TIRFM imaging.

### Mating experiments

For mating, 100 ml of vegetative cells were aerated in M medium under a light/dark cycle of 14:10 hours respectively for 4-5 days until a cell density of 2×10^6^ cells/ml was reached. The evening prior to the mating experiment, cells were transferred to 15 ml M-N medium and aerated overnight under constant light. In the morning, cells were transferred to 2 ml of 1/5 M-N supplemented with 10 mM HEPES and incubated for an additional 30 minutes, followed by mixing of gametes of opposite mating type and their incubation for 4-6 hours under light. Zygotes were observed using TIRFM. Progenies were obtained by following the mating protocol mentioned in (Liu et al., 2020).

### Isolation of cilia

To isolate cilia, cells were washed and concentrated in 10mM Hepes (pH 7.4), followed by resuspension in 10 ml of Hepes-Magnesium-Sucrose (HMS; 10 Mm Hepes, Ph 7.4, 5 mM MgSO_4,_ and simultaneous addition of 2.5 ml of dibucaine-HCl (25 mM in H_2_O; Sigma-Aldrich) to it and vigorous pipetting (reference) for deciliation. Next, 20 ml of 0.7 mM EGTA in HMS was added, and the cell bodies were removed by centrifugation (1,150 g, 3 min, 4°C; Sorvall Legend XTR, Thermo Fisher Scientific). A sucrose cushion (10 ML of 25% sucrose in HMS) was laid under the supernatant and centrifuged (1,700 × g, 4°C, 10 min) to remove the remaining cell bodies. The cilia in the upper phase were sedimented by centrifugation (27,000 g, 4°C, 20 min; Beckman Coulter, Avanti JXN-26) and resuspended in Hepes-Magnesium-EGTA-Potassium (HMEK; 30 mM Hepes, 5 mM MgSO_4_, 0.5 mM EGTA, and 25 mM KCl) supplemented with 1% protease inhibitor cocktail (PI, Sigma-Aldrich, P9599), and extracted for 20 min on ice with tx114 (1% final concentration). The axonemal and membrane + matrix fractions were separated by centrifugation (27,000 × *g*, 4°C, 15 min). Axonemes get separated as pellet and membrane +matrix remain in the supernatant. Supernatant is incubated at 30 C for 5 min which gives rise to a cloudy appearance and then centrifuged (1,700 g; 22 °C; 5 min). This leads to phase separation of the aqueous phase (matrix fraction) on top and detergent (membrane fraction) phase on bottom. The detergent phase is processed through methanol and chloroform precipitation to isolate the membrane.

### Antibodies and Western blotting

Anti-SIP and anti-MST1 antibodies were generated as follows: the coding region of *SIP* was amplified by PCR from *Chlamydomonas* cDNA using primers 1 and 2 (Table) and a ~500-bp long stretch encoded by exon 15 exon of *MST1* was amplified by PCR from a *Chlamydomonas* genomic DNA using primers 6 and 7 (Table 4). Both parts of *SIP* and *MST1* were cloned separately into the EcoRI site in the pMAL-cRI vector (downstream of the Maltose Binding Protein sequence). The fusion SIP and MST1 proteins were expressed in *E. coli,* purified using amylose resin according to the protocol provided by the manufacturer (New England Biolabs), and sent to Pocono Rabbit Farm and Laboratory for polyclonal antibody generation in rabbits. Sera containing the anti-SIP antibody were affinity-purified using SIP protein immobilized on PVDF membrane.

**Table 4).**
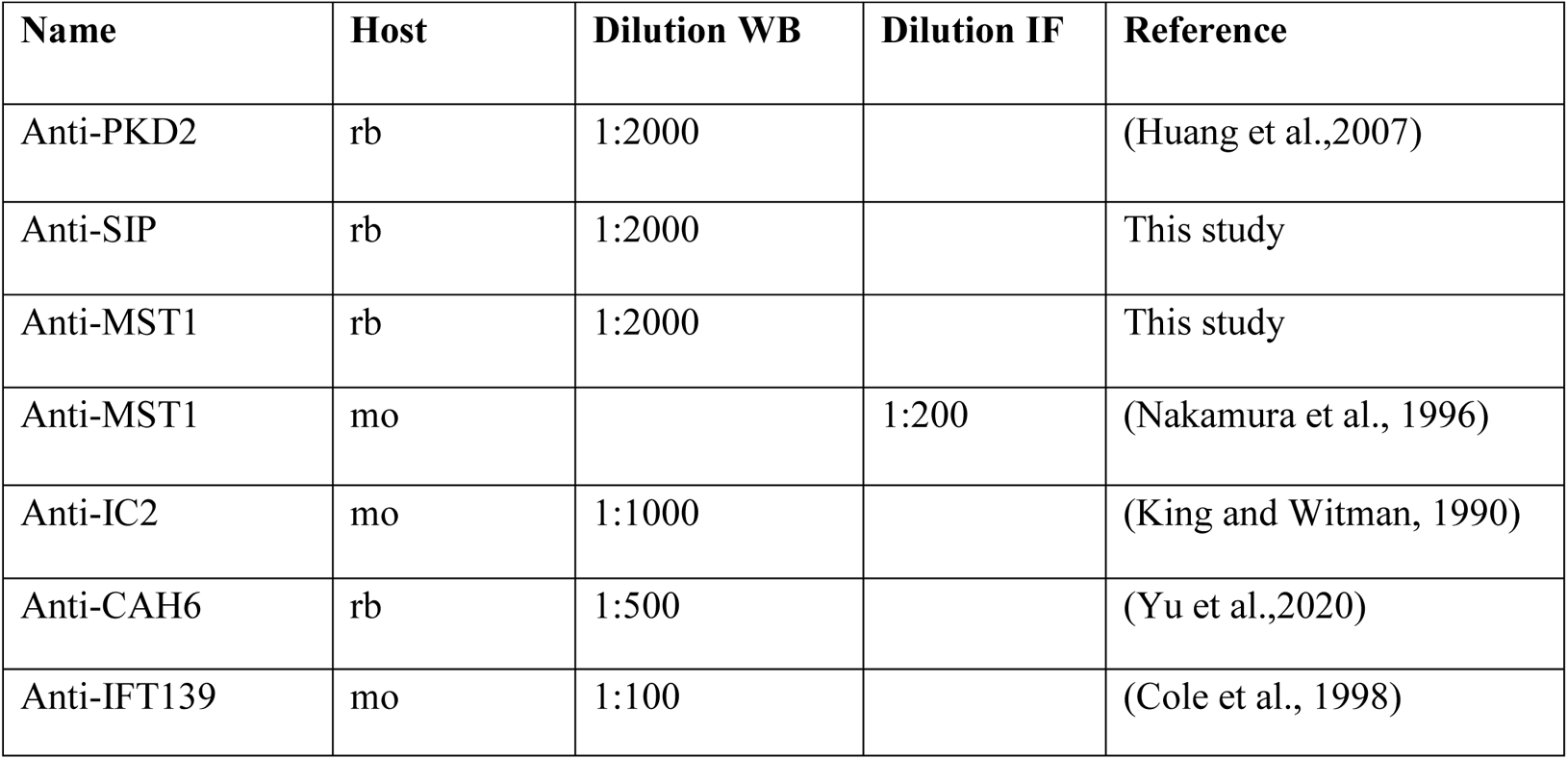
Antibodies used in this study.

Table 3) Primers used in this study
| Name | Host | Dilution WB | Dilution IF | Reference |
| --- | --- | --- | --- | --- |
| Anti-PKD2 | rb | 1:2000 |  | (Huang et al.,2007) |
| Anti-SIP | rb | 1:2000 |  | This study |
| Anti-MST1 | rb | 1:2000 |  | This study |
| Anti-MST1 | mo |  | 1:200 | (Nakamura et al., 1996) |
| Anti-IC2 | mo | 1:1000 |  | (King and Witman, 1990) |
| Anti-CAH6 | rb | 1:500 |  | (Yu et al.,2020) |
| Anti-IFT139 | mo | 1:100 |  | (Cole et al., 1998) |

Isolated cilia samples were incubated for 10 minutes at 95°C in Laemmli SDS sample buffer. Individual protein bands were separated by SDS-PAGE gels using Bio-Rad TGX precast gels and then transferred to PVDF membrane. Membranes were blocked in 3% BSA (Bovine Serum Albumin) and 3% fish gelatin followed by standard staining protocols, i.e., incubation in the diluted primary antibodies for overnight at 4°C with agitation (primary anti-bodies used in this study are listed in Table 4) and secondary antibodies (i.e., anti-mouse, 1:3000 and anti-rabbit IgG, 1:4000 conjugated to horseradish peroxidase; Invitrogen 31432/AB_228302 and 31460/AB_228341, respectively) at room temperature for ~60 minutes. For visualization, membranes were incubated in chemiluminescence substrate (SuperSignal West Pico PLUS or Atto; Thermo Fisher Scientific), and the images were captured using a Bio-Rad ChemiDoc MP imaging system and the Image Lab software. To quantify band intensities, we used Bio-Rad Image lab or the ROI Manager in ImageJ/Fiji.

### Immunoprecipitation

Cilia isolated from *pkd2* PKD2-mNG, *mst1 pkd2* PKD2-mNG and an untransformed control strain were resuspended in HMEK + protease inhibitor cocktail, lysed by addition of 1% NP-40 (final concentration), and 100 mM NaCl (final concentration). The axonemes were removed by centrifugation (27,000 *g*, 4°C, 15 min) and the supernatant was incubated with anti-NG nanobody agarose beads (Allele Biotechnology) for 1 hour at 4°C using a rotisserie. After loading the beads with protein, they were washed twice with HMEK containing 150mM NaCl. Proteins were eluted using 200 mM glycine, pH 2.5. The eluate, input, and flow-through were then analyzed using silver-stained gels (Silver Stain Plus Kit, Bio-Rad Laboratories) and the eluate was subjected to mass spectroscopy using an Orbitrap Elite system at the Proteomics and Mass Spectrometry Core Facility at the University of Georgia.

### Whole mount negative stain electron microscopy

For whole-mount negative stain, a formvar/carbon-coated 100 mesh electron microscope grid (FCF100-Cu-50, Electron Microscopy Sciences) was place on a drop of concentrated cells (~2 × 10^7^ cells/ml in water) on parafilm for 3 minutes. After removing excess cells using filter paper, the grid was put on a drop of 2% uranyl acetate in water for 1 to 2 minutes. Finally, the grid was washed with distilled water. Images were collected using a JEOL JEM1011 electron microscope.

### Swimming velocity and high-speed video

To measure the swimming velocity, cells were resuspended in fresh M medium, placed in a chambered plastic slide (14-377-259; Fisherbrand), and observed using an inverted light microscope (TMS; Nikon). Images were recorded using a MU500 camera (Amscope) and the associated Topview software at a fixed exposure time of 1 second. The length of the swimming trajectories (such as those shown in Fig. 6A) was analyzed using a newly developed “LengthAnalysis” plugin for ImageJ (National Institutes of Health)’; the plugin is described at https://github.com/Abha99/Length-Analysis-Tool. Details of the approach and a manual will be reported in the next version of the manuscript. Excel was used for statistical analysis and bar graphs and violin plots were prepared using GraphPad Prism.

### In vivo TIRF imaging

Samples for *in vivo* imaging were prepared as follows: at room temperature, 10 μl of cells were placed inside of a ring of petroleum jelly or vacuum grease onto a 24×60 mm No. 1.5 coverslip and allowed to settle for ~1-3 min. Then, a 22×22 mm No. 1.5 coverslip with 5 μl 10mM HEPES, 5mM EGTA, pH 7.4 was inverted onto the larger cover glass to form a sealed observation chamber. For gliding and secretion assay in Fig 4, EGTA was not included. For TIRF imaging, we used a Nikon Eclipse Ti-U inverted light microscope equipped with a 60x/1.49 NA objective lens and a 40 mW 488 nm diode lasers (Spectraphysics) (Lechtreck, 2013). Images were recorded at 10 fps using the iXon X3 DU897 EMCCD camera (Andor) and the Elements software package (Nikon). Still images mostly represent 10 frame walking averages. ImageJ (National Institutes of Health) with the Multiple Kymogram plug-in was used to analyze the recordings and generate kymograms. Kymograms, individual frames, and videos were cropped and adjusted for brightness and contrast in ImageJ and Photoshop CC 2018 (Adobe); Illustrator CC 2018 (Adobe) was used to assemble the figures.

### High-speed video analysis

For high-speed video analysis at 1000 fps, we used an inverted Eclipse Ti2 microscope (NIKON) equipped with a long distance DIC condenser and a 40x 0.95 Planapo objective. Images were recorded using an EoSens 3CL camera (Mikrotron) and an CORE2 DVR Express rapid storage device (IO Industries). Cells were concentrated and placed in an observation chamber. Recordings were exported in AVI format and analyzed using ImageJ.

## Abbreviations

DMT: doublet microtubule
NG: mNeonGreen
SIP: small interactor of PKD2

## Acknowledgements

This study was supported by a grant by the National Institutes of Health (R01GM139856 to K.L.). The content is solely the responsibility of the authors and does not necessarily represent the official views of the National Institutes of Health. B.M. and E.B.W. received CURO Research Assistantship for undergraduate researchers from the University of Georgia.

## Supplementary tables

**Table S1).**
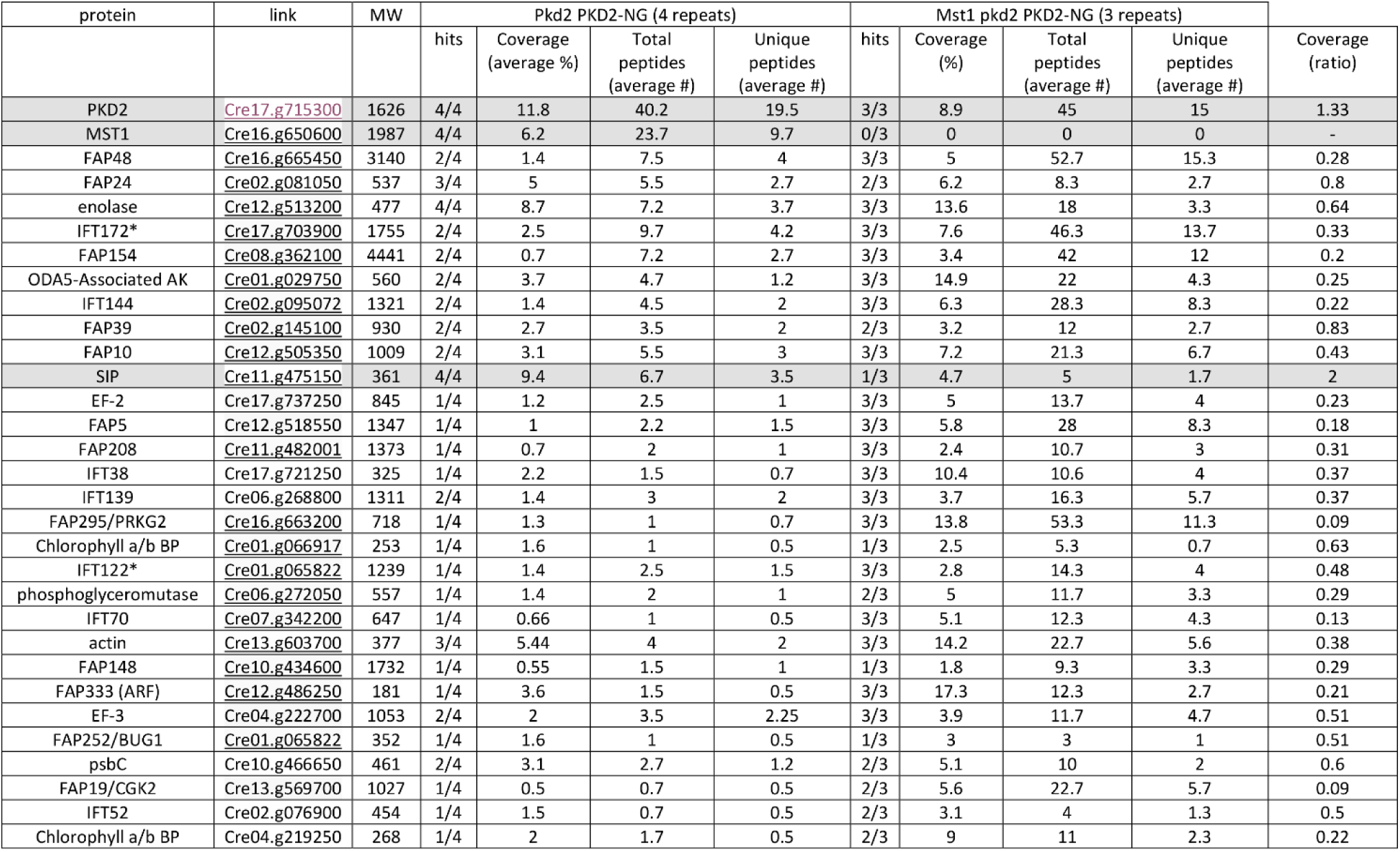
Mass spectrometry of PKD2-NG immune-isolates. The following proteins were found in all preparations, including those from the untagged control strain: In all preparations: FMG-1, FMG-1b, FAP33, FAP12, beta-tubulin, cobalamin-independent methionine synthase, eukaryotic translation elongation factor 1 alpha 1, putative blue light receptor, alpha-tubulin, IFT88, S-Adenosyl homocysteine hydrolase, IFT72/74, IFT81, IFT80, IFT57.

**Table S2).** Distribution and Conservation of SIP in green algae. NCBI Blastp and the *C. reinhardtii* sequences for SIP, PKD2, and MST1 were used search for related sequences.

|  | MINI-PKD2 (361) |  | MST1 (1987) |  | PKD2 (1645) |  |
| --- | --- | --- | --- | --- | --- | --- |
| Species | length | % identity<br>(% coverage) | length | % identity<br>(%coverage) | length | % identity<br>(% coverage) |
| *Chlamydomonas incerta* | 368 | 98.2 (93) | 1955 | 55.8 (89) | 1643 | 94.5 (99) |
| *Gonium pectorale* | 359 | 85.4 (91) | 868 | 52.8 (34) | 1629 | 76.5 (99) |
| *Volvox carteri* | 405 | 79.3 (91) | 835 | 51.5 (34) | 1606 | 67.3 (95) |
| *Trebuxia sp. A1-2* | 478 | 58.7 (73) | n.d. |  | 1618 | 48.9 (90) |
| *Chlorella ohadii* * | 456 | 51.2 (66) | n.d. |  | 1721 | 38.1 (73) |
| *Scenedesmus sp. NREL 46B-D3* * | 300 | 48.4 (78) | n.d. |  | 1582 | 43 (99) |
| *Dunaliella salina* | 360 | 37.5 (93) | n.d. |  | 1784 | 47.4 (95) |

## Supplementary figures

**Figure S1).**
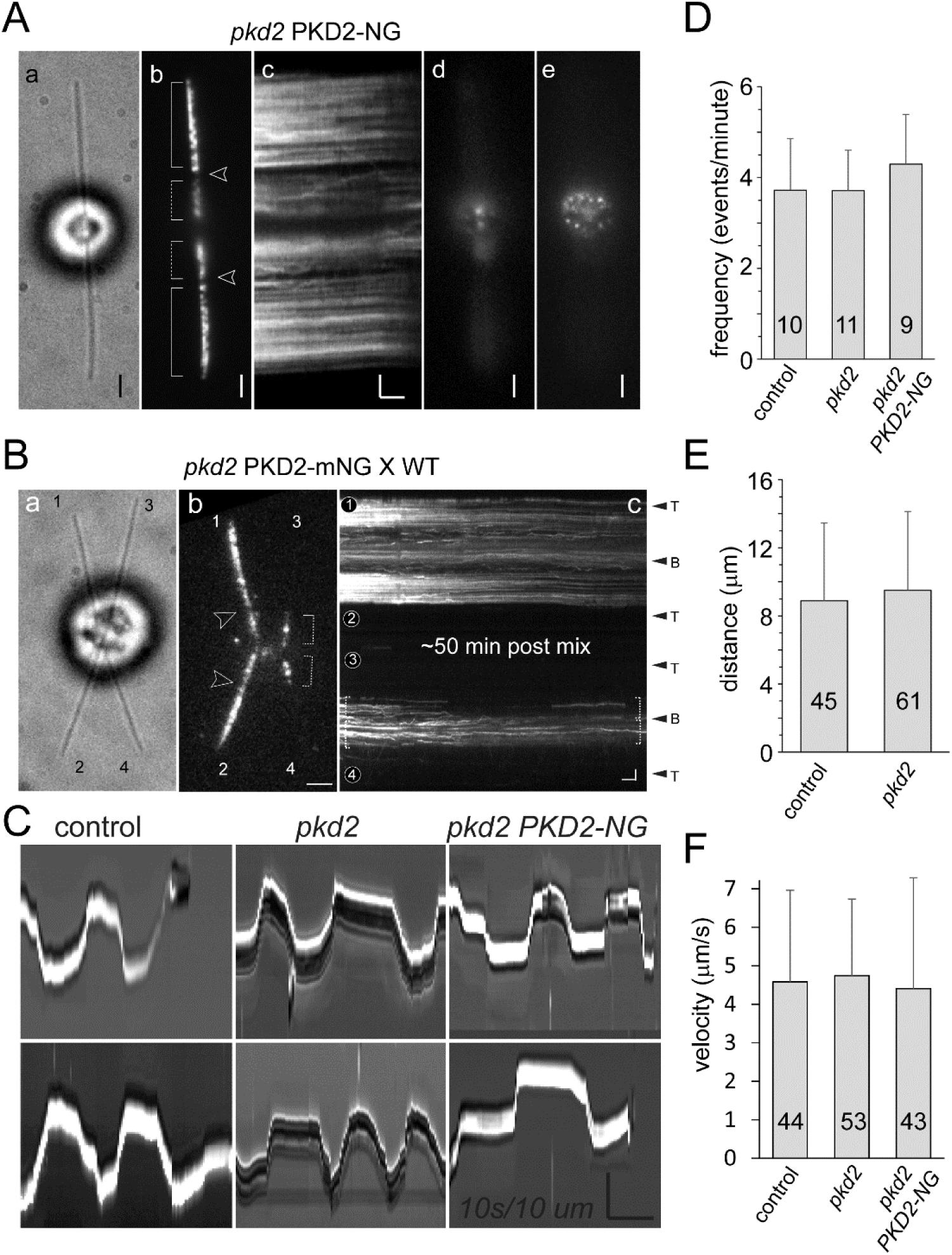
The pkd2 mutant displays normal gliding motility. A) Bright field (a), TIRF images (b, d, e) and corresponding kymogram (c) of a *pkd2 PKD2-NG* cell. B-e show different focal planes of the same cell showing the cilia (b) the basal body region (d) and the apical cell region (e). Note dotted distribution of PKD2-NG in the latter indicative for association with the sub-membranous microtubule cage of the cell body. Bars = 2s 2 µm. B) Bright field, TIRF image and kymogram of a quadriciliated *pkd2* PKD2-mNG × *pkd2* zygote. were scored by eye for the amount of PKD2-mNG in their cilia. Cilia derived from the *pkd2* PKD2-FP parent are labeled 1 and 2; those derived from the g1 control strain are labeled 3 and 4. The exclusion zone, proximal regions, and ciliary tips and bases are marked. Bars = 2 μm (in b for a and b) and 2 μm 2s (c). Note near absence of PKD2-NG form the distal segment of wild-type derived cilia. C) Kymograms of gliding control (g1) *pkd2*, and *pkd2* PKD2-mNG cells. Bars = 10 s 10μm. D-E) Analyses of gliding motility. Frequency (D), distance (E) and velocity (F) of gliding motion of the strains indicated. The number of events analyzed and the standard deviations are indicated. The differences between the strains were not significant.

**Figure S2).**
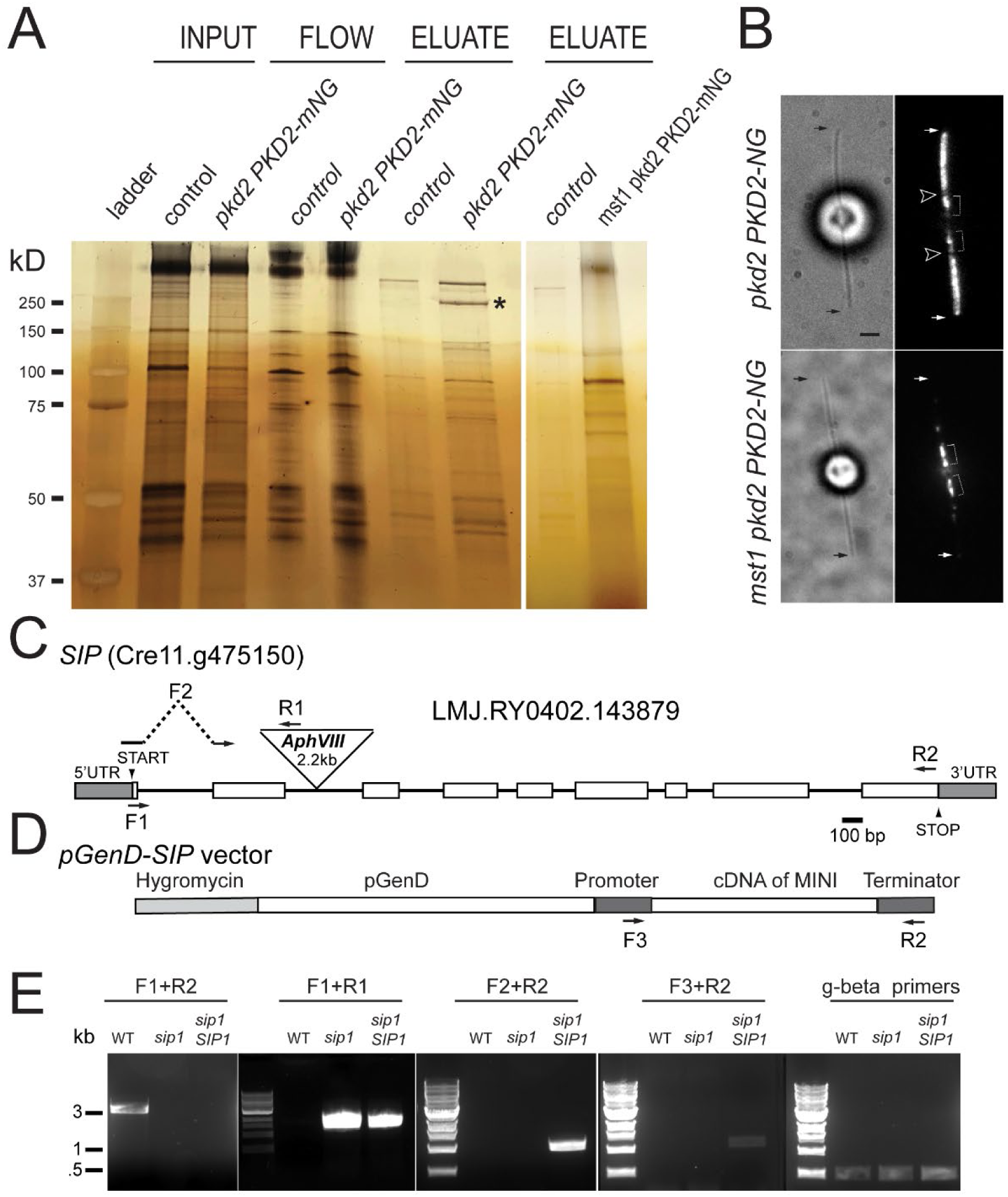
Identification of SIP as a PKD2 interacting protein. A) Silver stained gel analyzing the PKD2-mNG pull-down assays from a control strain (expressing endogenous PKD2), *pkd2* PKD2-mNG (null in endogenous PKD2 and expressing PKD2-mNG) and *mst1 pkd2* PKD2-mNG (null in MST1 and PKD2; expressing PKD2-mNG). The lanes are showing detergent extract input, flow through (FLOW), and eluates from anti-NG-nanobody sepharose trap. *, putative MST1 band. B) Bright field and TIRF images of pkd2 PKD2-NG and mst1 pkd2 PKD2-NG cells. The ciliary tips (small arrows) and the EZs (arrowheads) are marked. Bar = 2 um. C) Genomic map of the SIP/Cre11.g475150 gene. The positions of the *AphVIII* insertion and of the primers used to track the insertion (as shown in panel E) are indicated. D) Schematic presentation of the cDNA-based construct used to rescue the sip mutant. The positions of the primers used to track the transgene are indicated. E) Agarose gels of PCR reactions used to genotype control, *sip* mutant and *sip SIP* rescue cells.

**Figure S3).**
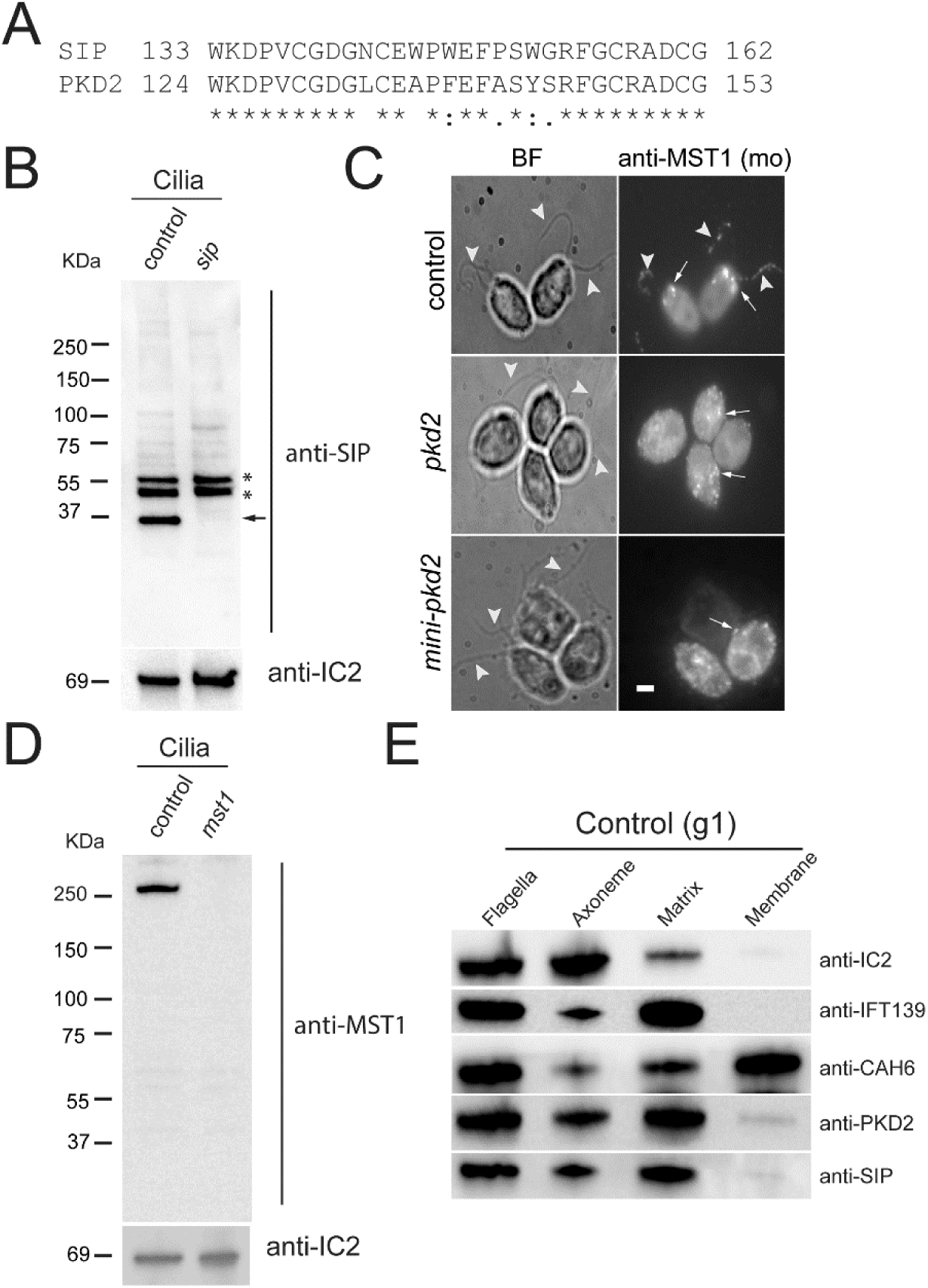
Antibodies to SIP and MST1. A) Conserved sequence stretches in *Chlamydomonas* PKD2 and SIP. B) Western blot of cilia isolated from control and the *sip* mutant and probed with anti-SIP. *, bands cross-reacting with anti-SIP. C) Immunofluorescence staining of control, pkd2 and sip mutant cells with monoclonal anti-MST1 (Nakamura et al., 1996) Arrowheads, cilia; the pool of MST1 in control cells and dispersed MST1 spots in *pkd2* and *sip* are marked by arrows. Bar = 2 µm. D) Western blot of cilia isolated from control and the *mst1* mutants and probed with polyclonal anti-MST1. In A and B, anti-IC2 was used as a loading control. E) Western blot analysis of cilia and ciliary fractions obtained by Triton X-114 phase partitioning from wild-type control strain (g1) using anti-SIP antibody to know SIP’s localization. As markers for the axonemal, membrane (i.e., detergent-soluble phase), and matrix (i.e., aqueous phase) fractions, we used antibodies to the ODA subunit IC2, the membrane-associated protein CAH6, and the IFT-B protein IFT139, respectively.

## References

Alfaro, G., Johansen, J., Dighe, S. A., Duamel, G., Kozminski, K. G. and Beh, C. T. (2011). The sterol-binding protein Kes1/Osh4p is a regulator of polarized exocytosis. Traffic 12, 1521–36.

Axelrod, D. (2003). Total internal reflection fluorescence microscopy in cell biology. Methods Enzymol 361, 1–33.

Bennett, H. W., Gustavsson, A. K., Bayas, C. A., Petrov, P. N., Mooney, N., Moerner, W. E. and Jackson, P. K. (2020). Novel fibrillar structure in the inversin compartment of primary cilia revealed by 3D single-molecule superresolution microscopy. Mol Biol Cell 31, 619–639.

Berman, S. A., Wilson, N. F., Haas, N. A. and Lefebvre, P. A. (2003). A novel MAP kinase regulates flagellar length in Chlamydomonas. Curr Biol 13, 1145–9.

Bloodgood, R. A. (1977). Motility occurring in association with the surface of the Chlamydomonas flagellum. J Cell Biol 75, 983–9.

Bloodgood, R. A. and Salomonsky, N. L. (1990). Calcium influx regulates antibody-induced glycoprotein movements within the Chlamydomonas flagellar membrane. J Cell Sci 96 (Pt 1), 27–33.

Bui, K. H., Yagi, T., Yamamoto, R., Kamiya, R. and Ishikawa, T. (2012). Polarity and asymmetry in the arrangement of dynein and related structures in the Chlamydomonas axoneme. J Cell Biol 198, 913–25.

Chung, J. J., Miki, K., Kim, D., Shim, S. H., Shi, H. F., Hwang, J. Y., Cai, X., Iseri, Y., Zhuang, X. and Clapham, D. E. (2017). CatSperzeta regulates the structural continuity of sperm Ca(2+) signaling domains and is required for normal fertility. Elife 6.

Chung, J. J., Shim, S. H., Everley, R. A., Gygi, S. P., Zhuang, X. and Clapham, D. E. (2014). Structurally distinct Ca(2+) signaling domains of sperm flagella orchestrate tyrosine phosphorylation and motility. Cell 157, 808–22.

DeCaen, P. G., Delling, M., Vien, T. N. and Clapham, D. E. (2013). Direct recording and molecular identification of the calcium channel of primary cilia. Nature 504, 315–8.

Dutcher, S. K. (1995). Flagellar assembly in two hundred and fifty easy-to-follow steps. Trends Genet 11, 398–404.

Dutcher, S. K. (2020). Asymmetries in the cilia of Chlamydomonas. Philos Trans R Soc Lond B Biol Sci 375, 20190153.

Dymek, E. E. and Smith, E. F. (2012). PF19 encodes the p60 catalytic subunit of katanin and is required for assembly of the flagellar central apparatus in Chlamydomonas. J Cell Sci 125, 3357–66.

Gadadhar, S., Hirschmugl, T. and Janke, C. (2023). The tubulin code in mammalian sperm development and function. Semin Cell Dev Biol 137, 26–37.

Geimer, S. and Melkonian, M. (2004). The ultrastructure of the Chlamydomonas reinhardtii basal apparatus: identification of an early marker of radial asymmetry inherent in the basal body. J Cell Sci 117, 2663–74.

Huang, K., Diener, D. R., Mitchell, A., Pazour, G. J., Witman, G. B. and Rosenbaum, J. L. (2007). Function and dynamics of PKD2 in Chlamydomonas reinhardtii flagella. J Cell Biol 179, 501–14.

Isom, L. L. (2001). Sodium channel beta subunits: anything but auxiliary. Neuroscientist 7, 42–54.

Kottgen, M., Hofherr, A., Li, W., Chu, K., Cook, S., Montell, C. and Watnick, T. (2011). Drosophila sperm swim backwards in the female reproductive tract and are activated via TRPP2 ion channels. PLoS One 6, e20031.

Kozminski, K. G., Beech, P. L. and Rosenbaum, J. L. (1995). The Chlamydomonas kinesin-like protein FLA10 is involved in motility associated with the flagellar membrane. J Cell Biol 131, 1517–27.

Laib, J. A., Marin, J. A., Bloodgood, R. A. and Guilford, W. H. (2009). The reciprocal coordination and mechanics of molecular motors in living cells. Proc Natl Acad Sci U S A 106, 3190–5.

Lechtreck, K. F. (2013). In vivo Imaging of IFT in Chlamydomonas Flagella. Methods Enzymol 524, 265–84.

Lechtreck, K. F. (2016). Methods for Studying Movement of Molecules Within Cilia. Methods Mol Biol 1454, 83–96.

Lee, S., Tan, H. Y., Geneva, II, Kruglov, A. and Calvert, P. D. (2018). Actin filaments partition primary cilia membranes into distinct fluid corrals. J Cell Biol 217, 2831–2849.

Lefebvre, P. A., Nordstrom, S. A., Moulder, J. E. and Rosenbaum, J. L. (1978). Flagellar elongation and shortening in Chlamydomonas. IV. Effects of flagellar detachment, regeneration, and resorption on the induction of flagellar protein synthesis. J Cell Biol 78, 8–27.

Li, X., Patena, W., Fauser, F., Jinkerson, R. E., Saroussi, S., Meyer, M. T., Ivanova, N., Robertson, J. M., Yue, R., Zhang, R. et al. (2019). A genome-wide algal mutant library and functional screen identifies genes required for eukaryotic photosynthesis. Nat Genet 51, 627–635.

Liu, P., Lou, X., Wingfield, J. L., Lin, J., Nicastro, D. and Lechtreck, K. (2020). Chlamydomonas PKD2 organizes mastigonemes, hair-like glycoprotein polymers on cilia. J Cell Biol 219.

Mahjoub, M. R., Montpetit, B., Zhao, L., Finst, R. J., Goh, B., Kim, A. C. and Quarmby, L. M. (2002). The FA2 gene of Chlamydomonas encodes a NIMA family kinase with roles in cell cycle progression and microtubule severing during deflagellation. J Cell Sci 115, 1759–68.

Matsuzaki, O., Bakin, R. E., Cai, X., Menco, B. P. and Ronnett, G. V. (1999). Localization of the olfactory cyclic nucleotide-gated channel subunit 1 in normal, embryonic and regenerating olfactory epithelium. Neuroscience 94, 131–40.

Mochizuki, T., Saijoh, Y., Tsuchiya, K., Shirayoshi, Y., Takai, S., Taya, C., Yonekawa, H., Yamada, K., Nihei, H., Nakatsuji, N. et al. (1998). Cloning of inv, a gene that controls left/right asymmetry and kidney development. Nature 395, 177–81.

Nakamura, S., Takino, H. and Kojima, M. K. (1987). Effect of Lithium on Flagellar Length in Chlamydomonas-Reinhardtii. Cell Structure and Function 12, 369–374.

Nakamura, S., Tanaka, G., Maeda, T., Kamiya, R., Matsunaga, T. and Nikaido, O. (1996). Assembly and function of Chlamydomonas flagellar mastigonemes as probed with a monoclonal antibody. J Cell Sci 109 (Pt 1), 57–62.

Pazour, G. J., Agrin, N., Leszyk, J. and Witman, G. B. (2005). Proteomic analysis of a eukaryotic cilium. J Cell Biol 170, 103–13.

Pazour, G. J., Sineshchekov, O. A. and Witman, G. B. (1995). Mutational analysis of the phototransduction pathway of Chlamydomonas reinhardtii. J Cell Biol 131, 427–40.

Shiba, D., Yamaoka, Y., Hagiwara, H., Takamatsu, T., Hamada, H. and Yokoyama, T. (2009). Localization of Inv in a distinctive intraciliary compartment requires the C-terminal ninein-homolog-containing region. J Cell Sci 122, 44–54.

Shih, S. M., Engel, B. D., Kocabas, F., Bilyard, T., Gennerich, A., Marshall, W. F. and Yildiz, A. (2013). Intraflagellar transport drives flagellar surface motility. Elife 2, e00744.

Silflow, C. D. and Lefebvre, P. A. (2001). Assembly and motility of eukaryotic cilia and flagella. Lessons from Chlamydomonas reinhardtii. Plant Physiol 127, 1500–7.

Singh, A. P. and Rajender, S. (2015). CatSper channel, sperm function and male fertility. Reprod Biomed Online 30, 28–38.

Tam, L. W., Ranum, P. T. and Lefebvre, P. A. (2013). CDKL5 regulates flagellar length and localizes to the base of the flagella in Chlamydomonas. Mol Biol Cell 24, 588–600.

van der Burght, S. N., Rademakers, S., Johnson, J. L., Li, C., Kremers, G. J., Houtsmuller, A. B., Leroux, M. R. and Jansen, G. (2020). Ciliary Tip Signaling Compartment Is Formed and Maintained by Intraflagellar Transport. Curr Biol 30, 4299–4306 e5.

Walther, Z., Vashishtha, M. and Hall, J. L. (1994). The Chlamydomonas FLA10 gene encodes a novel kinesin-homologous protein. J Cell Biol 126, 175–88.

Wang, J., Nikonorova, I. A., Gu, A., Sternberg, P. W. and Barr, M. M. (2020). Release and targeting of polycystin-2-carrying ciliary extracellular vesicles. Curr Biol 30, R755–R756.

Watnick, T. J., Jin, Y., Matunis, E., Kernan, M. J. and Montell, C. (2003). A flagellar polycystin-2 homolog required for male fertility in Drosophila. Curr Biol 13, 2179–84.

Wilson, N. F. and Lefebvre, P. A. (2004). Regulation of flagellar assembly by glycogen synthase kinase 3 in Chlamydomonas reinhardtii. Eukaryot Cell 3, 1307–19.

Xiang, W., Zur Lage, P., Newton, F. G., Qiu, G. and Jarman, A. P. (2022). The dynamics of protein localisation to restricted zones within Drosophila mechanosensory cilia. Sci Rep 12, 13338.

Yagi, T., Uematsu, K., Liu, Z. and Kamiya, R. (2009). Identification of dyneins that localize exclusively to the proximal portion of Chlamydomonas flagella. J Cell Sci 122, 1306–14.

Zamora, I., Feldman, J. L. and Marshall, W. F. (2004). PCR-based assay for mating type and diploidy in Chlamydomonas. Biotechniques 37, 534–6.

Zhang, W., Cheng, L. E., Kittelmann, M., Li, J., Petkovic, M., Cheng, T., Jin, P., Guo, Z., Gopfert, M. C., Jan, L. Y. et al. (2015). Ankyrin Repeats Convey Force to Gate the NOMPC Mechanotransduction Channel. Cell 162, 1391–403.

Zhao, Y., Wang, H., Wiesehoefer, C., Shah, N. B., Reetz, E., Hwang, J. Y., Huang, X., Wang, T. E., Lishko, P. V., Davies, K. M. et al. (2022). 3D structure and in situ arrangements of CatSper channel in the sperm flagellum. Nat Commun 13, 3439.

